# Dissecting the molecular triggers of early and late long-term potentiation

**DOI:** 10.64898/2026.04.09.717511

**Authors:** Rui Wang, Michaela Schweizer, Kristina Ponimaskine, Christian Schulze, Christine E. Gee, Thomas G. Oertner

## Abstract

The brain stores information by changing the strength of its synapses, a process that has at least two phases: Late long-term potentiation (L-LTP) is thought to result from the consolidation of early LTP (E-LTP), just as long-term memory requires the prior establishment of short-term memory. Recently, inhibitory avoidance experiments under CaMKII inhibition have challenged this notion, demonstrating long-term fear memory without measurable short-term memory. Here we use optogenetic activation and inhibition of CaMKII during induction of spike-timing-dependent potentiation (tLTP) to dissect the signaling pathways. While CaMKII activation in CA1 neurons was sufficient to induce E-LTP, growth of the postsynaptic density and spine neck expansion, we found that CaMKII-induced LTP does not give rise to L-LTP. Conversely, inhibition of CaMKII during tLTP induction prevented E-LTP, but FOS and L-LTP were still expressed, driven by CaMKK and PKMζ. Thus, both long-term memory and L-LTP form in the absence of CaMKII activation.

## Introduction

Long-term potentiation (LTP) is a persistent increase in synaptic strength that has long been considered a cellular correlate of learning and memory. LTP can be divided into at least two temporally and mechanistically distinct phases: early LTP (E-LTP), which lasts from minutes to hours, depends on receptor insertion and post-translational modifications, and can be reversed; and late LTP (L-LTP), which can persist for days, requires gene transcription and protein synthesis (*1*). This mirrors the distinction between short-term and long-term memory, implying that long-lasting plasticity sequentially consolidates from transient forms. The activation of calcium/calmodulin-dependent protein kinase II (CaMKII) is critical for the induction of E-LTP. CaMKII is rapidly recruited to the postsynaptic density following calcium influx through NMDA receptors (*2*). Activated CaMKII then phosphorylates AMPA receptor subunits and synaptic scaffolding proteins. This leads to the insertion and stabilization of AMPA receptors (*3*, *4*) and the enlargement of dendritic spines (*5*). Thus, activation of CaMKII is considered both necessary and sufficient for the induction of E-LTP. The formation of CaMKII/NMDA receptor complexes through binding to GluN2B is critical for long-term memory (*6–8*). The molecular requirements for L-LTP and its relationship to E-LTP remain less clear. Classical models suggested that L-LTP results from the consolidation of E-LTP through protein synthesis and gene transcription triggered by the initial synaptic potentiation (*9*, *10*). However, recent behavioral studies have challenged this sequential model. Inhibitory avoidance experiments performed under CaMKII inhibition revealed that long-term fear memory can form in the absence of short-term memory (*11*). This finding raises the possibility that the signaling cascades responsible for long-term memory formation may be activated independently of the classical CaMKII-dependent E-LTP pathway (*12*).

To dissect the temporal and mechanistic contributions of CaMKII signaling to E-LTP and L-LTP at Schaffer collateral synapses, we combined optogenetic manipulation of CaMKII (*4*, *13*) with an optogenetic spike-timing-dependent plasticity (STDP) protocol (*14*). Optogenetic spike-timing-dependent potentiation (tLTP) can be induced under sterile conditions inside a cell culture incubator or during patch-clamp recordings, which allowed us to assess the strength of synaptic connections either immediately (E-LTP) or three days later (L-LTP). By precisely activating or inhibiting CaMKII in defined neuronal populations, we found that L-LTP can emerge independently of E-LTP and identified alternative pathways supporting its delayed expression. Our findings reveal that while optical activation of CaMKII is sufficient to induce E-LTP and synapse remodeling, the strength of these artificially potentiated synapses slowly declines. Conversely, inhibition of CaMKII during tLTP induction prevented E-LTP, but still allowed for the development of L-LTP and expression of immediate early genes. Post-induction pharmacology revealed CaMKK (*15*) and PKMζ (*16*, *17*) as key mediators of L-LTP. The requirement for CaMKK activity hours after the induction protocol suggests that L-LTP in CA1 requires ongoing network activity, reminiscent of the active consolidation of hippocampal memories during sleep (*12*, *18*).

## Results

### Structural synaptic plasticity induced by optogenetic activation of paCaMKII

To monitor structural changes at many synapses simultaneously, we used a customized two-photon microscope with stage incubator (37°C, 5% CO_2_) optimized for long-term imaging of hippocampal slice cultures (see Methods). CA1 pyramidal cells were individually electroporated with a mix of three plasmids: The monobody Xph20–eGFP–CCR5TC (*19*) to label postsynaptic densities (PSDs), a long Stokes shift orange-red fluorescent protein (LSSmOrange) (*20*) to label the cytoplasm, and photoactivatable CaMKII (paCaMKII) (*4*) to trigger synaptic plasticity (Fig. 1A). Chronic 3D imaging was performed on oblique dendrites in *stratum radiatum* (Fig. 1B). Green fluorescent PSDs were automatically identified (*21*) (Fig. 1C) and tracked over 16 hours. Spine volume was estimated from the red fluorescence intensity at every PSD location. Following global photoactivation of paCaMKII, PSDs and spine head volumes increased rapidly (Fig. 1D, Fig. S1, Fig. S2), consistent with prior single-spine activation studies (*4*). Experiments without light stimulation, or light stimulation of neurons transfected with paCaMKII-SD, an inactive control plasmid, did not display rapid changes in PSD size or spine volume and were thus pooled into one unified control group. In line with previous reports (*22–26*), the distributions of spine volume and PSD size were right-skewed with a long tail towards high values (Fig. 1F). To determine whether the growth induced by CaMKII activation depended on the initial size of the synapse, we calculated the growth factor of each PSD and spine head (change in intensity 2 - 5 h after optical stimulation normalized to baseline) and plotted it against each synapse’s baseline intensity. For statistical analysis, synapses were binned into quartiles according to their baseline intensity (Fig. 1F). Relative to non-activated controls, both PSD and spine head volumes increased significantly across all four size quartiles. The largest relative effect occurred in the smallest synapses: Small PSDs increased by ∼50% and small spines by ∼30%. In contrast, the largest synapses showed more modest changes (large PSDs ∼22%, large spines ∼11%). In the next hours, PSDs continued to grow in the stimulated neurons (Fig. 1F), indicating that a single brief paCaMKII activation (100 s) initiated a prolonged growth program that affected both spine head volume and PSD size.

**Figure 1:**
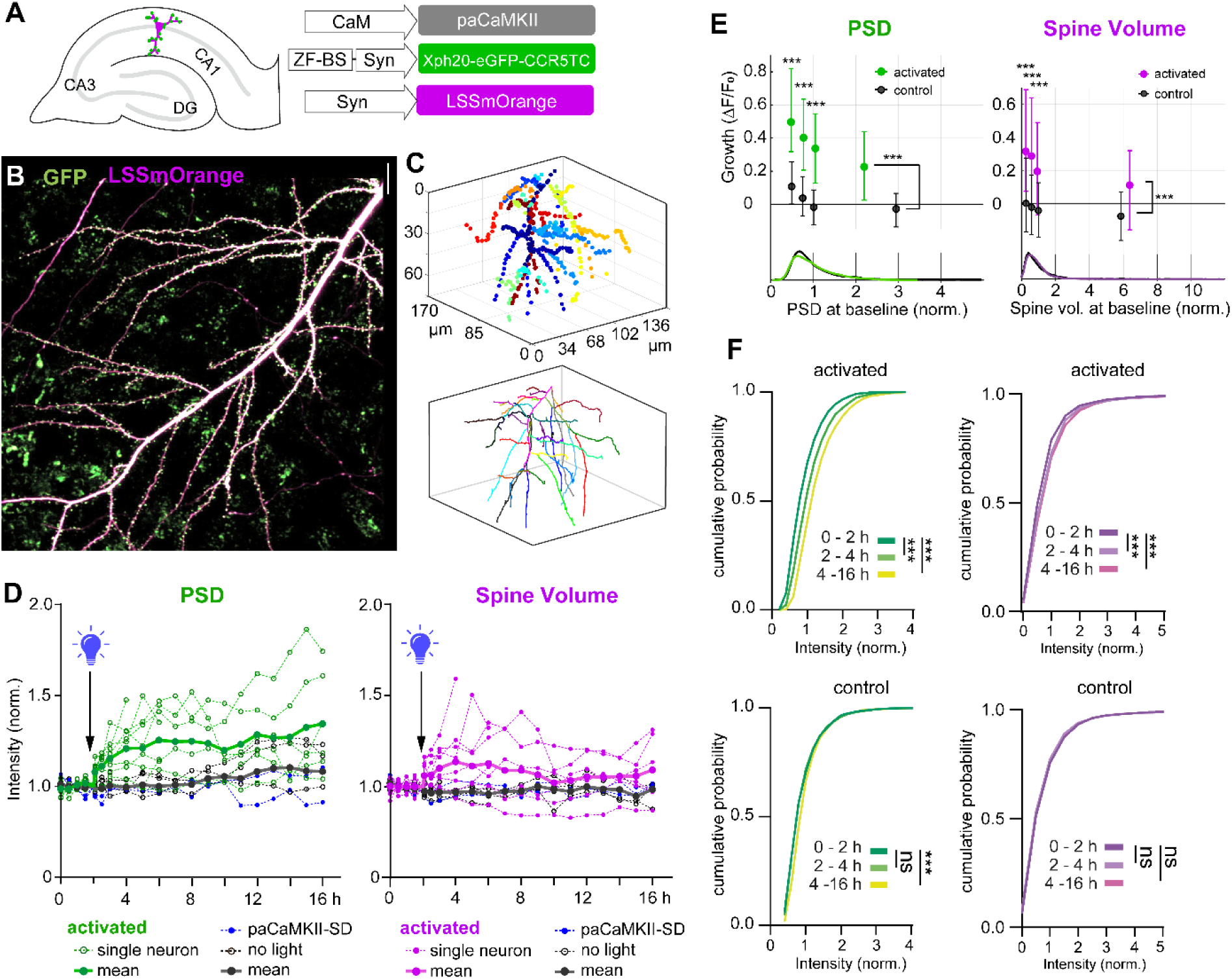
Optical activation of CaMKII induces PSD and spine growth. (**A**) Strategy of triple transfection and synapse imaging in CA1 pyramidal neurons. (**B**) Spiny dendrites in *stratum radiatum* (magenta: cytoplasm; green: monobody against PSD95). (**C**) Automatically detected synapses in 3D, color-coded by dendritic branch. (**D**) PSD (green) and spine head growth (magenta) induced by paCaMKII activation at t = 2 h (blue lightbulb, n = 9 neurons). Control experiments without paCaMKII activation are shown in gray. Control group consists of 5 paCaMKII neurons that were not light-stimulated (open circles), and 2 paCaMKII-SD neurons that were light-stimulated (blue dots). Dotted lines show average of all PSDs/spines from individual cultures, solid lines show average of averages. n = 3947 and 5799 PSDs in activated and control conditions. (**E**) Growth analysis in size groups (quartiles, size distribution at baseline shown below) 2 - 5 h after optical stimulation (mean ± SEM). paCaMKII activation induced significant growth of PSDs (green) and spine volumes (magenta) in all size groups compared to controls (black). Activated: n = 608, 608, 609, 608 synapses (PSDs, spines) from 4 slice cultures. Control (black), n = 738, 738, 739, 738 synapses from 5 slice cultures. Data plotted as median ± quartiles. Mann-Whitney test, ***p < 0.001. (**F**) Cumulative distribution of individual PSDs (green) and spine volumes (magenta) at baseline (0 to 2 h), early (2 to 4 h) and late (4 to 16 h) after the activation of paCaMKII, in activated and control conditions. N = 6, 5 (slice cultures); n = 2948, 2953 (PSDs). Kolmogorov-Smirnov test, ***p < 0.0001.

### Spine neck remodeling following paCaMKII activation reduces synaptic compartmentalization

Two-photon microscopy allowed us to monitor the total size of each PSD over time, but the spatial resolution of light microscopy is not sufficient to determine the shape of the spine head and its neck. Repeated glutamate uncaging in Mg^2+^-free solution has been shown to shorten and widen the necks of mushroom spines (*27*). Therefore, we investigated whether CaMKII activation alone, bypassing the activation of NMDA receptors and calcium influx, would be sufficient to drive similar structural changes. We co-transfected CA1 pyramidal cells with paCaMKII and dAPEX2, a peroxidase optimized for electron microscopy (EM) reconstruction of fine neuronal processes (*28*, *29*). We fixed the transfected slice cultures 10 min after optical activation of paCaMKII and performed electron tomography of labeled synapses (Fig. S3). Following 3D reconstruction that was performed blind to the stimulation conditions, we observed enlarged spine heads and significant remodeling of spine neck geometry. Spine necks were significantly thicker and shorter after paCaMKII activation (Fig. 2A, B), indicating rapid effects of active CaMKII on the actin cytoskeleton. The size of the PSD was not different from controls at this early time point. Monobody labeling of PSD-95 had already increased by 8% (Fig. 1D), but it may take more time to assemble all proteins that contribute to the eponymous enhanced electron density.

**Figure 2:**
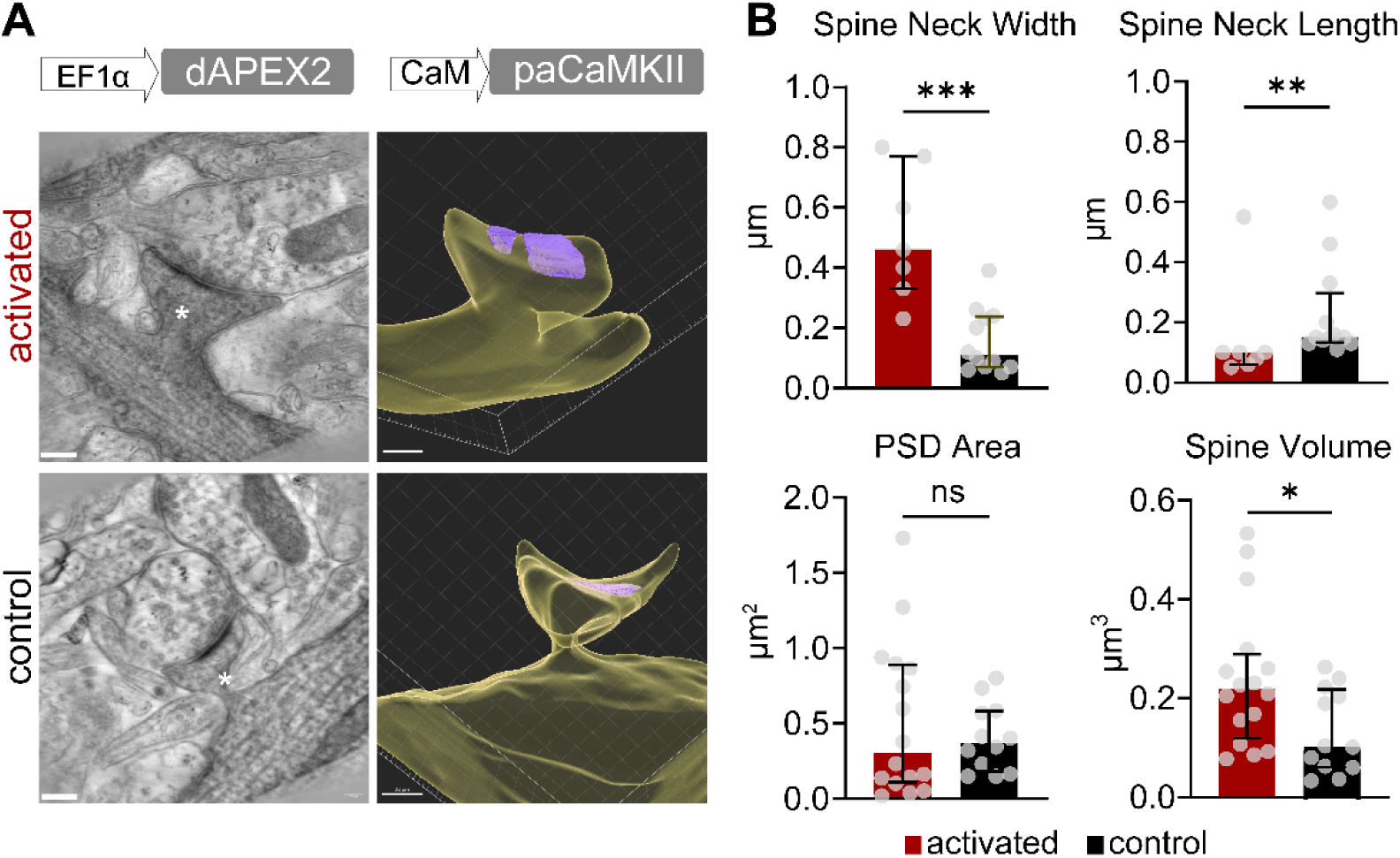
Optical activation of CaMKII remodels spine neck geometry. (**A**) dAPEX2 tagging to identify paCaMKII-transfected spines in EM tomograms. Example images and corresponding 3D reconstructions show labeled spines with or without paCaMKII activation. Yellow: spine head and connected dendrites; magenta: PSD. Scale bars: no light: 200 nm; activated: 300 nm. (**B**) Analysis of spine structure (paCaMKII activation: red, n = 7 reconstructed spines from 2 slice cultures vs. control: black, n = 12 reconstructed spines from 2 slice cultures) and PSD area (paCaMKII activation: red, n = 16 reconstructed spines from 2 slice cultures vs. control: black, n = 12 reconstructed spines from 2 slice cultures). Data plotted as median ± quartiles. Mann-Whitney test, ***p < 0.001, **p < 0.01, *p < 0.05, ns: p > 0.05.

Changes in the geometry of the spine neck could affect the compartmentalization of proteins in the spine head. It has been proposed that widening and shortening of spine necks could contribute to LTP by reducing the electrical resistance between synapse and dendrite (*27*, *30*). On the other hand, cofilin-actin plugs form in the spine neck after strong stimulation and may restrict diffusion (*31*, *32*). To measure the diffusional coupling between spine and dendrite before and after paCaMKII activation, we performed two-photon Fluorescence Recovery After Photobleaching (FRAP, Fig. 3A) experiments. As expected, the red cytoplasmic fluorescence rapidly recovered, indicating diffusional coupling between spine and dendrite, while bleached PSD95-monobody fluorescence (green) did not recover within the observation window (6 s, Fig. 3B), confirming that the intrabodies were firmly anchored to the postsynaptic density. The recovery time constant of cytoplasmic fluorescence (τ, determined by single exponential fit) was heterogeneous between spines, with values ranging from less than 100 ms to over 2 seconds, reflecting the diversity of spine head volumes and neck shapes (*27*, *33*). After paCaMKII activation, a small but significant reduction in τ was observed (Fig. 3C and D), while the photoactivation light pulse had no systematic effect on cells expressing the non-functional control plasmid paCaMKII-SD. The FRAP data provide functional evidence that CaMKII-induced neck remodeling effectively lowers the diffusional barrier between spine head and parent dendrite.

**Figure 3:**
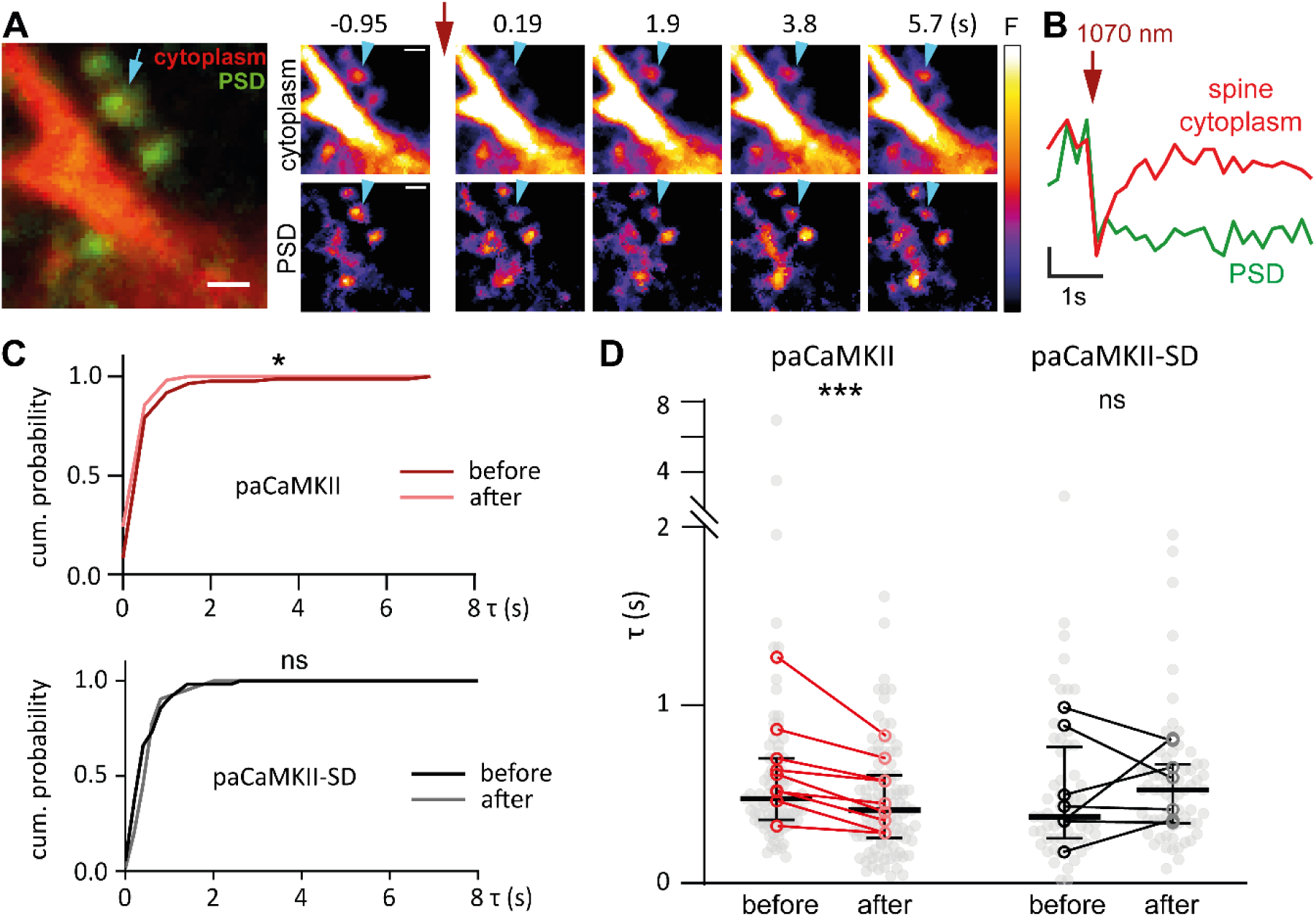
Optical activation of CaMKII reduces synaptic compartmentalization. (**A**) Two-photon FRAP experiments on individual spines in CA1. Neurons expressed paCaMKII, Xph20-eGFP-CCR5TC (green) and tdimer2 (red). After baseline imaging (represented by the t = −0.95 s image), a short bleaching pulse was delivered (t = 0, red flash) to the target spine (cyan arrow) and the recovery of fluorescence was monitored. Scale bars: 1 μm. (**B**) Spine head fluorescence recovered (red) while monobody fluorescence (green) did not. Scale bars: 20% dF/F_0_, 1 s. (**C**) The cumulative distribution of FRAP time constants (τ) changed 5-10 min after paCaMKII activation. Spines on neurons expressing paCaMKII-SD (super dark, light-insensitive) were used as controls (gray). Kolmogorov-Smirnov test, n = 86, 104, 56, 63 spines. *p < 0.05. (**D**) Effects of light stimulation on synaptic compartmentalization in individual slice cultures (same data as C). Light grey dots show τ values of individual spines, black lines show median ± quartiles. Red circles show means of individual slice cultures before and after light stimulation (n = 9, 9, 7, 7 cultures, two-tailed Wilcoxon signed-rank test, pairwise comparisons performed on culture means, ***p < 0.001. Total number of spines analyzed: 86, 104, 56, 63.

### Functional LTP induced by paCaMKII activation lacks long-term stability

Biochemical assays and intracellular delivery of purified, activated CaMKII have been used to demonstrate that CaMKII activation is sufficient to induce E-LTP (20-30 min) (*34*). Two-photon activation of paCaMKII at single synapses induced functional E-LTP (25-35 min) and enlargement of spine heads that lasted at least 4 hours (*4*). Our chronic imaging experiments confirmed the spine head expansion and in addition, demonstrated PSD enlargement that was still significant 14 h after plasticity induction. Next, we wanted to evaluate E- and L-LTP using whole-cell recordings from paCaMKII-expressing CA1 pyramidal cells. For these experiments, we combined single-cell paCaMKII transfection in CA1 with channelrhodopsin transduction (ChrimsonR) in CA3 via local AVV injection. To test the function of this approach, we performed whole-cell recordings to measure red light-evoked EPSCs (Fig. 4A). Low-intensity violet light (405 nm, 0.1 mW/mm^2^, 100 seconds) activated paCaMKII and rapidly induced LTP that lasted for at least 25 minutes (Fig. 4B, C). Control CA1 neurons expressing a non-functional plasmid (paCaMKII-SD) did not show potentiation when subjected to the same optical stimulation. The paCaMKII-induced LTP was independent of NMDA receptor function (Fig. S4), in line with previous findings that the intracellular activation of paCaMKII does not evoke Ca^2+^ influx (*4*). To test the input strength over multi-day timescales, we used again optical stimulation inside the incubator to activate paCaMKII under sterile conditions (*14*). To test for L-LTP, the strength of CA3 inputs to paCaMKII-activated CA1 neurons (EPSC slope) was measured 1 and 2 days after paCaMKII activation by whole-cell patch-clamp recordings from paCaMKII-expressing neurons and normalized to the response of several non-transfected neighbors (Fig. S5) (*14*). No input strengthening was observed 1 or 2 days after paCaMKII activation (Fig. 4D, Fig. S6). Thus, one-time paCaMKII activation did produce early, but not long-lasting LTP.

**Figure 4:**
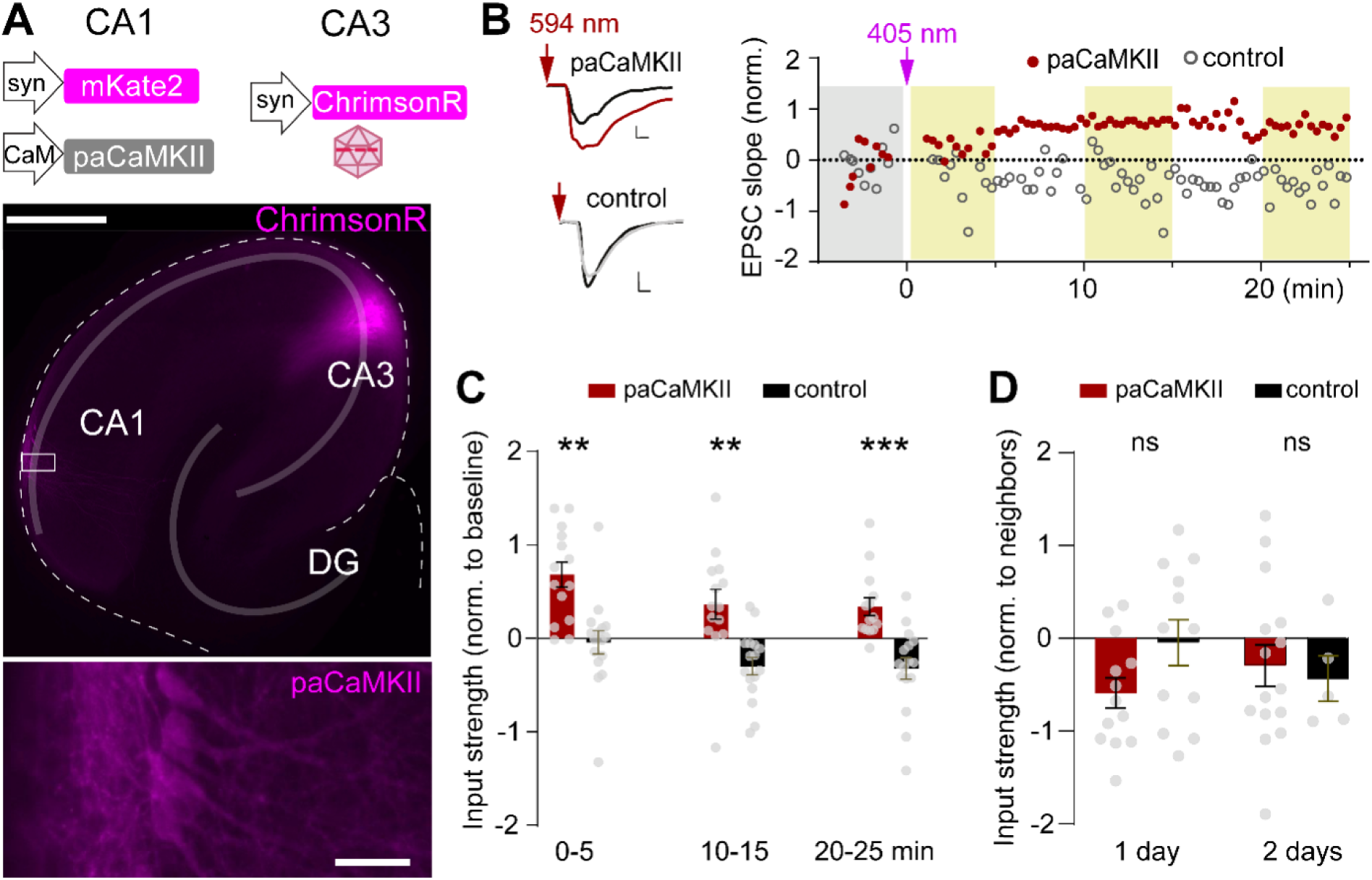
CaMKII activation is sufficient to induce early, but not late LTP. (**A**) Strategy of transfection and the representative image showing CA3 cells virally transduced with ChrimsonR and CA1 cells electroporated with paCaMKII (white box, enlarged). (**B**) Red light pulses activate ChrimsonR, evoking EPSCs in a patch-clamped CA1 neuron. A single violet light pulse (405 nm) at t = 0 activates paCaMKII, changing the slope of EPSCs. Traces show average EPSC during baseline (black) and 20-25 min after paCaMKII activation (red). Control neuron expresses paCaMKII-SD (black, gray). Scale bars: 100 pA, 5 ms. (**C**) Activation of paCaMKII induces significant LTP (n = 14 cultures) compared to control neurons expressing paCaMKII-SD (control, n = 16 cultures). Two-way ANOVA followed by Dunnett’s multiple comparison test, ***p < 0.001. Gray points show individual experiments (cells), bars show mean ± SEM. (**D**) Results from in-incubator stimulation experiments, assessed 1 or 2 days after activation of paCaMKII. No difference between neurons expressing paCaMKII (n = 13, 15) and paCaMKII-SD (control, n = 12, 5). Unpaired t-test, p = 0.072, 0.73. Gray points show individual experiments (EPSC slope in transfected neuron normalized to 3 neighboring non-transfected neurons, see Fig. S5). Bars show mean ± SEM.

It has been proposed that neurons that undergo LTP express immediate early genes, such as FOS and JUN (*35*). These proteins form the AP-1 transcription factor, which triggers the expression of various plasticity-related proteins and increases neuronal excitability (*36*). However, one hour after paCaMKII activation, we did not detect significant FOS expression in paCaMKII-expressing CA1 neurons (Fig. 5). As a positive control, we elevated the extracellular K^+^ concentration, which triggered strong FOS expression in paCaMKII-expressing CA1 neurons and their neighbors. This provides an important clue as to why optogenetic CaMKII activation was not sufficient to induce long-lasting LTP: If cell-wide CaMKII activity is not sufficient to trigger the genomic response characteristic for L-LTP, other calcium-sensing pathways must be involved.

**Figure 5:**
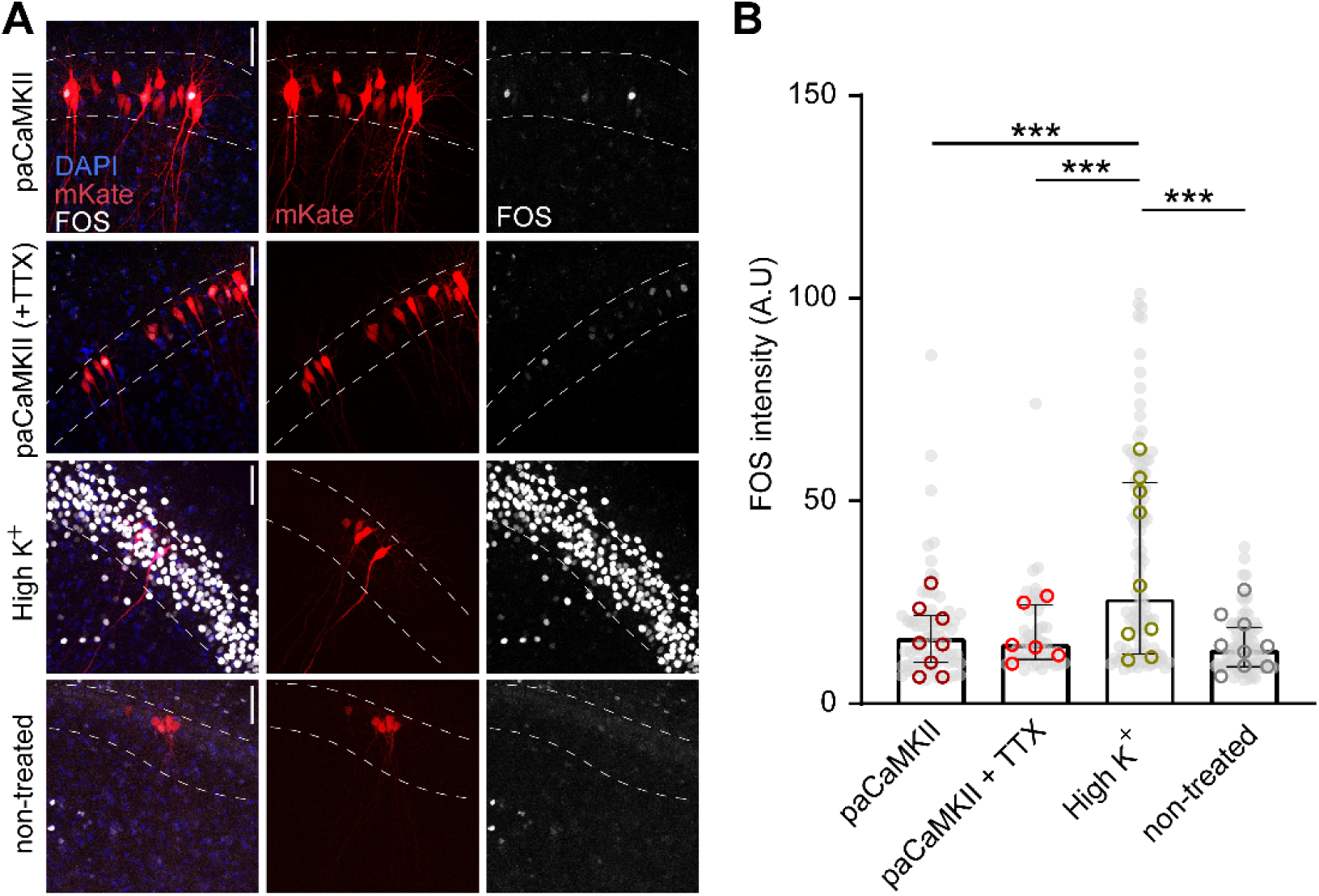
CAMKII activation does not induce significant increase of FOS expression. (**A**) Immunostaining 1 h after light stimulation to activate paCaMKII (representative images). In the upper two rows, red neurons express paCaMKII. In the lower two rows (positive and negative control), red neurons express just mKate2. Anti-FOS immunoreactivity is shown in white. Dashed lines indicate the cell body layer. Scale bars: 50 μm. (**B**) Evaluation of FOS immunoreactivity in red fluorescent neurons. FOS expression after paCaMKII activation was not different from negative (non-treated) controls. Kruskal-Wallis test followed by Dunn’s multiple comparisons, ***p < 0.001, n = 71, 52, 104, 76 cells (gray dots), bars show median ± quartiles. Open circles: Replicates (8, 6, 9, 9 slice cultures). A.U., arbitrary units.

### Optical inhibition of CaMKII reveals the dissociation of early-LTP and late-LTP

We next asked whether the CaMKII activity is required for spike-timing-dependent plasticity (STDP) (*37*). We previously developed an all-optical method to induce timing-dependent LTP (tLTP) in hippocampal slice cultures (*14*). Two different channelrhodopsins, ChrimsonR and CheRiff, are used to control the spike timing in pre- and postsynaptic neurons. ChrimsonR, activated by red light, is expressed in CA3 neurons and CheRiff, activated by violet light, is expressed in CA1 (Fig. 6A, B). Repeatedly pairing presynaptic action potentials with short postsynaptic bursts (10 ms delay, ‘causal pairing’) results in stable tLTP that can be detected 3 days later (*14*). Here we combine this optical pairing protocol with paAIP2, a photoactivatable inhibitor of CaMKII (*13*) with an activation spectrum similar to CheRiff. Consequently, in neurons that express both paAIP2 and CheRiff, endogenous CaMKII is inhibited during optical pairing (Fig. 6A). As a positive control, we used neurons expressing only CheRiff. The first set of tLTP experiments was performed while recording EPSCs in the postsynaptic CA1 neuron (whole-cell patch-clamp). CA3, which was outside the field of view of the objective, was illuminated through the condenser. Twenty minutes after optical pairing, tLTP was induced in CheRiff-expressing neurons, but not in neurons expressing CheRiff and paAIP2 (Fig. 6C, D), suggesting that CaMKII activation was essential for the induction of tLTP. Recent studies suggested that CaMKII silencing can depress basal synaptic transmission (*38*, *39*), arguing that the lack of LTP in the CaMKII-inhibited condition is caused by a run-down of synaptic transmission. To test this possibility, we applied the same causal pairing protocol to CA1 neurons expressing only paAIP2 (no CheRiff). In each slice culture, paAIP2-only and non-transfected (NT) neurons were patched sequentially, and the same presynaptic optical stimulation pulses were applied to record baseline transmission. After recording each baseline, a pairing protocol was performed (in the absence of postsynaptic CheRiff). The baseline slope of each pair of paAIP2 and non-transfected cells was identical (Fig. 6E), suggesting that expression of paAIP2 did not reduce the input strength. More importantly, after optical inhibition of CaMKII, no reduction in input strength was observed in the paAIP2 group (Fig. 6E, Fig. S7). These results show that acute optogenetic inhibition of CaMKII does not affect synaptic transmission, but blocks the induction of synaptic plasticity. Therefore, CaMKII activation induced by coincident spikes in the pre- and the postsynaptic neuron is indeed critical for the early phase of tLTP in the hippocampus.

**Figure 6:**
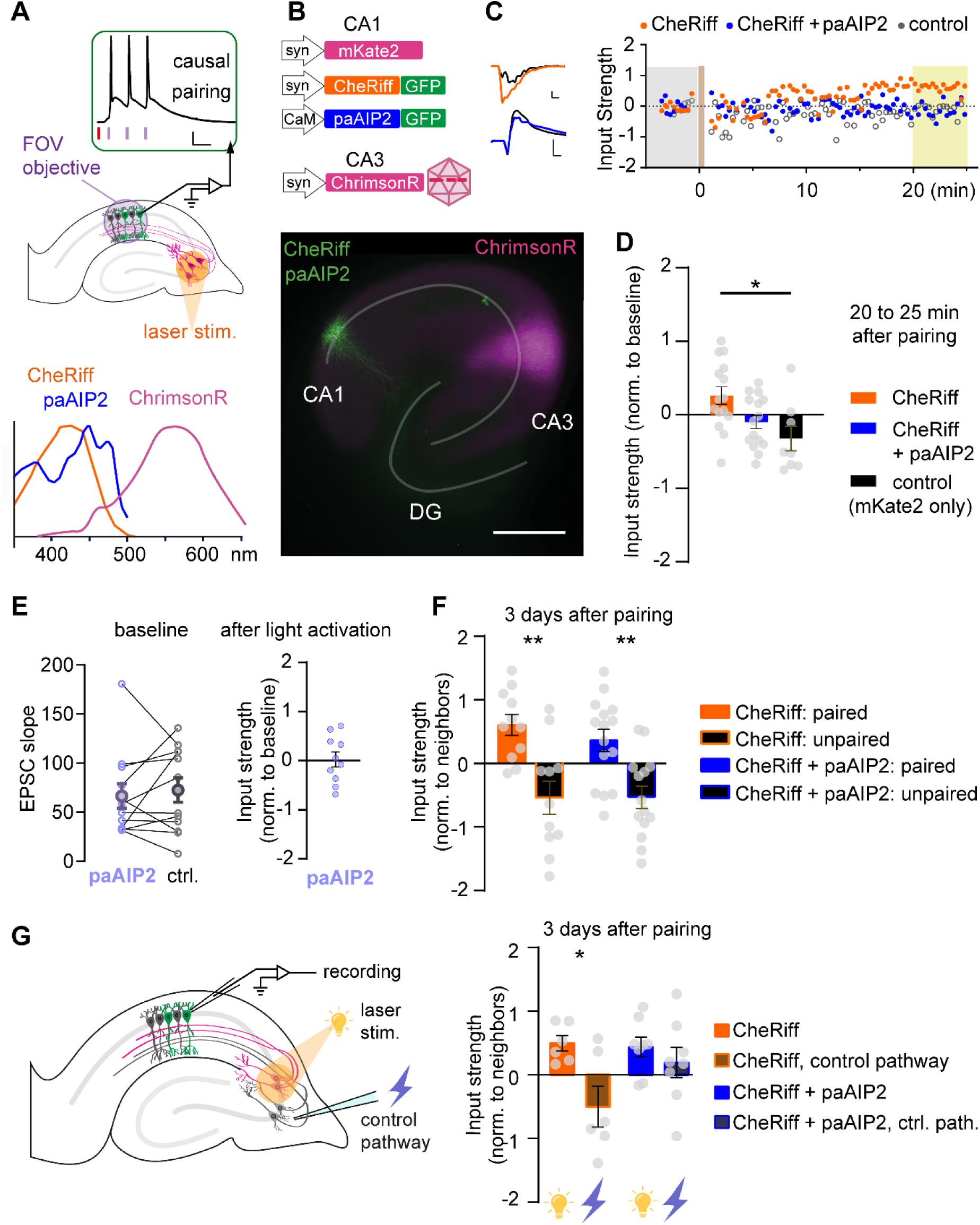
CaMKII activity is critical for the early, but not for the late phase of synaptic plasticity. (**A**) Optical pairing protocol. Red light activation of CA3 neurons is followed by 3 violet light pulses to spike CA1 neurons. Below: Spectral sensitivity of paAIP2, CheRiff and ChrimsonR (Adapted from Murakoshi et al., 2017 & Nagasawa et al., 2023). (**B**) CA1 neurons were transfected with CheRiff-GFP and mKate2, or with CheRiff-GFP, mKate2 and paAIP2, or just with mKate2. CA3 neurons were virally transduced with ChrimsonR. (**C**) Light-induced EPSCs at baseline (black) and 20-25 minutes after tLTP induction. The arrow indicates the time of red-light flashes used to spike the CA3 neurons. Example whole-cell patch clamp recordings of synaptic strength alteration upon tLTP stimulation (yellow-violet bar at t = 0). paAIP2 activation blocks the inductions of tLTP. Control neurons received light stimulation, but expressed only mKate2. Scale bars: 100pA, 5 ms. (**D**) Input strength 25 minutes after pairing protocol, normalized to baseline. CheRiff-expressing neurons, but not neurons co-expressing paAIP2, are significantly different from the control group. NT, CheRiff, CheRiff +paAIP2: n = 8, 15, 15 slice cultures; Data plotted as mean ± SEM. Ordinary one-way ANOVA, followed by Tukey’s multiple comparisons. (**E**) Left: average EPSC slopes of sequentially recorded paAIP2 and NT neurons (ctrl.). Paired comparisons were performed between the two cells recorded from the same slice, optical stimulation was applied to each slice. Open circles: individual recordings; filled circles: mean ± SEM. Paired t-test. NT, paAIP2: n = 12, 12 (cells). Right: Optogenetic inhibition of CaMKII activity for 25 min does not affect synaptic strength. n = 10 paAIP2-expressing cells. (**F**) Optogenetic inhibition of CaMKII activity does not block late tLTP. Unpaired controls expressed identical constructs, but did not receive light stimulation. Left to right: n = 11, 11, 15, 14 slice cultures. Data plotted as mean ± SEM. Unpaired t-test, **p < 0.01. **(G)** Synapse specificity of tLTP. Three days after induction of tLTP, EPSCs were evoked by optical stimulation (potentiated pathway) or by electrical stimulation in CA3 (control pathway). Orange bars show outcome with CaMKII signaling intact, blue bars show the outcome when CaMKII signaling was blocked during induction via paAIP2 activation (mean ± SEM). Left to right: n = 6, 6, 8, 8 slice cultures. Unpaired t-test, p = 0.044.

To address L-LTP, we repeated the tLTP experiments using in-incubator stimulation and assessed the results 3 days after plasticity induction in CheRiff-only and CheRiff + paAIP2 expressing neurons. The optogenetic tLTP induction protocol incl. paAIP2 activation was identical for E-LTP and L-LTP experiments (matched light intensities and wavelengths). As expected, optically induced L-LTP was detected in CheRiff-only expressing neurons, compared to their non-transfected neighbors (Fig. 6F, Fig. S6). Surprisingly, inputs to neurons expressing CheRiff and paAIP2 were also potentiated, even though we could not detect any E-LTP in this group. This suggests that signaling pathways driving a slowly developing, long-lasting form of LTP are still activated by the tLTP protocol in the absence of initial CaMKII-dependent potentiation. Thus, L-LTP is not simply a continuation or stabilization of E-LTP, but relies on completely different induction pathways that do not require CaMKII activity and take considerable time to fully express.

The synaptic tagging and capture (STC) hypothesis proposes that late-phase LTP remains synapse-specific because molecular tags established during early LTP enable potentiated synapses to selectively capture plasticity-related proteins produced during the late phase (*1*). NMDAR-anchored, persistently active CaMKII is a candidate molecular component of the synaptic tag (*8*). To test whether tLTP induced under conditions of CaMKII inhibition remains synapse-specific, we next examined two independent input pathways. As before, optogenetic reactivation of CA3 neurons was used to assess the strength of synapses that were directly activated during the STDP induction protocol. These synapses were potentiated 3 days after STDP, irrespective of whether CaMKII was inhibited during induction (Fig. 6G). In parallel, a stimulation electrode was placed in CA3 to activate a distinct set of Schaffer collateral inputs that had not been activated during the STDP protocol. With CaMKII function intact, this control pathway was depressed 3 days after STDP, demonstrating the strong synapse specificity of tLTP. In contrast, when CaMKII was inhibited during induction, the difference between the potentiated and control pathways was abolished. These findings indicate that CaMKII activation during early LTP is required for the synapse-specific expression of late LTP, providing experimental support for a central prediction of the STC hypothesis (*40*).

The canonical pathway to L-LTP requires CREB phosphorylation, activation of immediate early genes, and production of plasticity-related proteins. One hour after optical pairing, we observed robust FOS expression in CheRiff-expressing CA1 neurons, but not in their neighbors (Fig. 7A). Surprisingly, FOS expression was even stronger in CheRiff + paAIP2 neurons, reaching levels similar to the positive control (K^+^) (Fig. 7B, C). Again, FOS expression was highly specific to the paired neurons; no FOS was detected in neighboring neurons that also received synaptic stimulation from CA3. These findings provide a molecular explanation for the delayed emergence of L-LTP. The pairing protocol triggers a parallel, CaMKII-independent signaling pathway that relays information to the nucleus, initiating the gene expression programs required for L-LTP even under conditions that prevent E-LTP.

**Figure 7:**
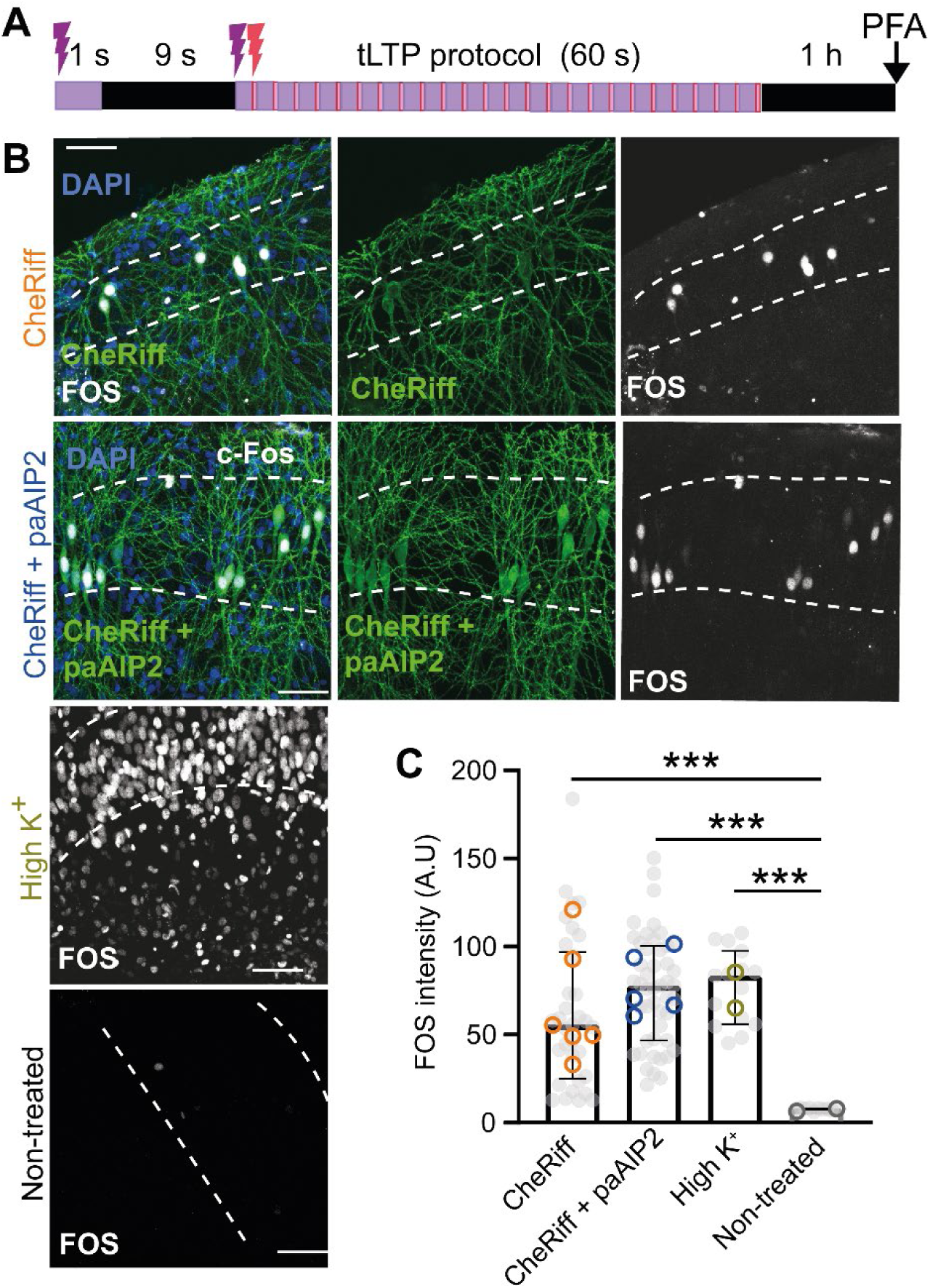
The activation of CaMKII is dispensable for tLTP-induced FOS expression. (**A**) Experimental protocol. Causal pairing stimulation (violet-red light flashes for 60 seconds) was applied in the incubator, with CaMKII activity blocked 9 seconds ahead of time, by 405 nm pulses for 1 s. After the stimulation the slice culture was kept in the incubator for one hour, then fixed for immunohistochemistry staining. (**B**) Immunostaining 1 h after optical pairing. Cell body layer is indicated by the dash lines. Anti-FOS immunoreactivity is shown in white. Scale bar:50 μm. (**C**) Evaluation of FOS immunoreactivity. With (‘CheRiff’) or without (‘CheRiff + paAIP2’) CaMKII activity, FOS expression after optical pairing activation was not different from positive (high K+) controls. Kruskal-Wallis test followed by Dunn’s multiple comparisons test, ***p < 0.001, n = 33, 47, 15, 10 cells (grey dots); bars show median ± quartiles. Open circles: Replicates (6, 5, 2, 2 slice cultures). A.U., arbitrary units.

### CaMKK, but not CaMKII, governs late-phase LTP

Our data highlights the distinct molecular mechanisms of early and late long-term plasticity. To identify pathways critical for L-LTP, we used a post-induction blockade protocol (Fig. 8A). Ca^2+^/calmodulin-dependent protein kinase kinase (CaMKK) is considered essential for expression of plasticity-related proteins and L-LTP (*15*). Inhibition of CaMKK activity starting 3 hours after inducing tLTP (STO-609) blocked CaMKII-dependent and CaMKII-independent L-tLTP (Fig. 8B). STO-609 applied before inducing tLTP also prevented FOS expression in the CheRiff-expressing neurons (Fig. S8), confirming the key role of this enzyme. PKMζ, an atypical isoform of PKC, serves as a critical plasticity-related protein that is selectively recruited to and retained by potentiated synapses (*16*). Application of the PKMζ peptide inhibitor ZIP (ζ-inhibitor peptide) also blocked L-tLTP in both groups (Fig. 8B). In control experiments with scrambled protein (scr-ZIP), L-tLTP was not affected. Prolonged incubation (2 days) with either inhibitor did not alter neuronal viability or intrinsic electrophysiological properties (Fig. S9), ruling out neurotoxicity (*41*, *42*) as a confounding factor. Collectively, our results delineate a fundamental functional and molecular divergence between the early and late phases of plasticity (Fig. 8C). We demonstrate that while optogenetic CaMKII activation is sufficient to drive rapid structural remodeling, it fails to engage the signaling cascades required for long-term stability. The so-called consolidation of synaptic memory relies on a parallel, CaMKII-independent signaling axis, with CaMKK and PKMζ required to couple coincident synaptic activation to the nuclear gene expression programs necessary for the slow-onset development and maintenance of late-LTP.

**Figure 8:**
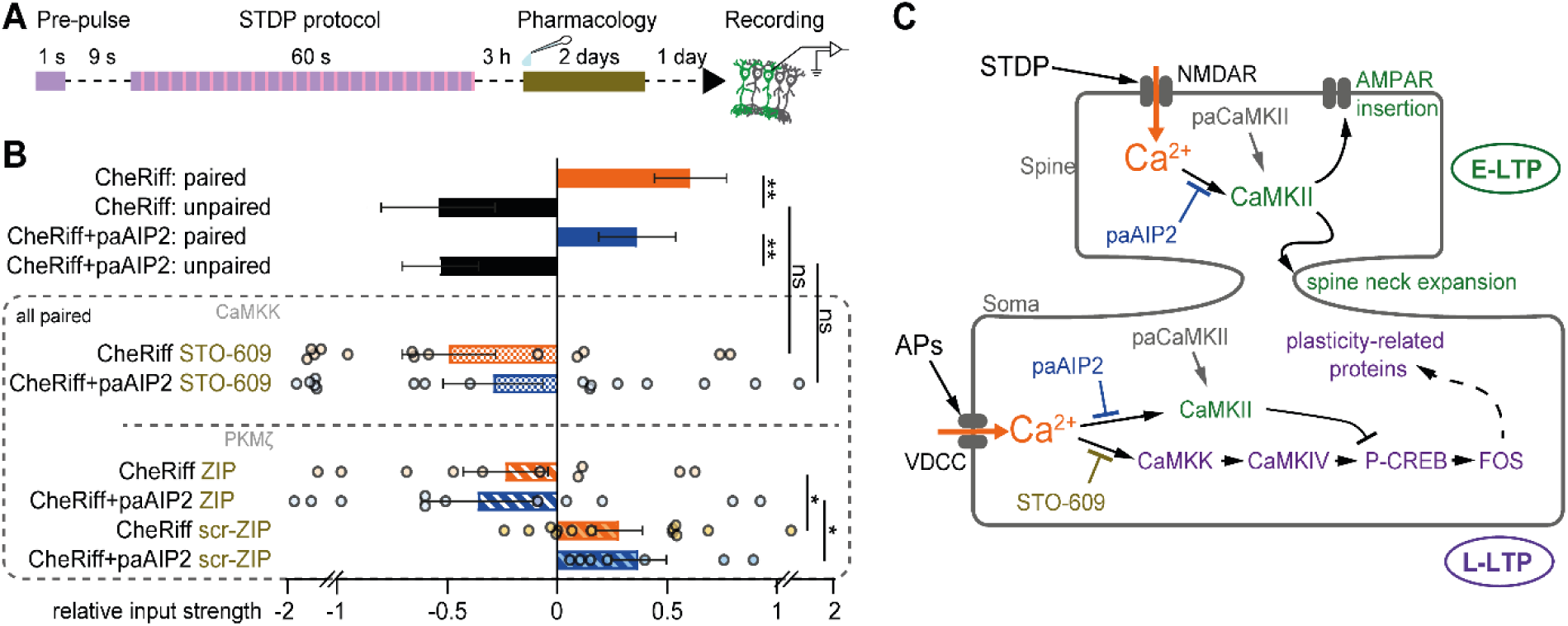
Late LTP requires CaMKK, but not CaMKII activity. (**A**) Experimental protocol. Causal pairing stimulation (violet-red light flashes for 60 s) was applied in the incubator. In paAIP experiments, a pre-pulse (405 nm) was given to activate the inhibitor. After the stimulation, the slice culture was kept in the incubator for three hours before the application of pharmacological inhibitors. After 2 days of treatment, slices were moved back to drug-free medium for 1 more day before electrophysiological recordings. (**B**) Comparisons of input strength 3 days after pairing shows that the CaMKK antagonist STO-609 (5 µM) and the PKMζ blocking peptide ZIP (1 µM) prevented CaMKII-dependent and CaMKII-independent late tLTP. Unpaired t-test, **p < 0.01, *p < 0.05. Data plotted as mean ± SEM. Top 4 bars are re-plotted of Fig. 6F. n = 11, 11, 15, 14, 13, 16, 10, 11, 13, 7 cells. (**C**) The tLTP protocol activates NMDARs and voltage-dependent calcium channels (VDCCs), the resulting Ca^2+^ influx triggers two distinct signaling cascades: E-LTP (green) requires CaMKII activation; L-LTP (magenta) requires CaMKK-CaMKIV-CREB signaling in the soma/nucleus to initiate the synthesis of plasticity-related proteins.

## Discussion

It is generally assumed that L-LTP forms the basis for memory and requires the prior formation of CaMKII-dependent E-LTP (*1*, *43*). A recent study challenged this sequential model by showing that CaMKII inhibition abolishes short-term fear memory and synaptic potentiation in the amygdala, but unexpectedly, long-term memory would still develop under these conditions (*11*). Our results reveal distinct signaling mechanisms underlying the rapid induction of E-LTP and the delayed expression of L-LTP. Consistent with previous single-spine studies (*4*), we show that acute activation of paCaMKII is sufficient to induce rapid synaptic structural enlargement of postsynaptic densities, spine neck constriction, and strengthening of synaptic transmission. However, this potentiation was transient and largely dissipated within 1 day. The simultaneous potentiation of many synapses may have engaged homeostatic downscaling (*44*), thereby masking the subsequent expression of L-LTP. Importantly, despite activation of CaMKII across all synapses of a given postsynaptic neuron, this manipulation failed to engage the genomic machinery associated with stable synaptic strengthening, as reflected by the absence of FOS induction. As previously demonstrated (*4*), activation of paCaMKII does not increase intracellular calcium levels, providing a potential explanation for its inability to engage the signaling pathways required for L-LTP. Conversely, optical inhibition of postsynaptic CaMKII during tLTP induction abolished E-LTP while leaving L-LTP intact. Together, these findings argue against a model in which L-LTP simply represents the maintenance or consolidation of E-LTP. Instead, they support a model in which L-LTP is initiated through a distinct signaling pathway that operates in parallel with rapid, CaMKII-dependent E-LTP and engages a genomic program required for persistent synaptic strengthening (Fig. 8C).

Two methodological advances enabled the isolation and identification of parallel signaling pathways: tLTP induction under sterile conditions, achieved without patching the neurons; and the precise timing and specificity of optogenetic CaMKII inhibition and activation. Controlling the spike timing of two populations of neurons inside the cell culture incubator enabled us to uncouple tLTP induction from connection strength measurement, extending the tLTP observation window from minutes to days (*14*, *29*). This extended temporal window was essential for identifying two kinases, CaMKK and PKMζ, which are required in the postsynaptic neuron for late tLTP. The ability to transiently and selectively inhibit and activate CaMKII in the postsynaptic neuron excludes direct effects on the presynaptic compartment or non-neuronal cells (*45*). This makes it relatively straightforward to explain the results in terms of postsynaptic mechanisms (Fig. 8C).

A critical question arises: How is the synapse successfully potentiated when CaMKII activation, the canonical master regulator, is inhibited? Our data identify CaMKK as an essential effector that bypasses the inhibition of CaMKII activity and autonomously initiates L-LTP. Ca^2+^/calmodulin binds to CaMKK and relieves its autoinhibition mechanism. However, CaMKK cannot sustain enzymatic activity in the absence of Ca^2+^. Therefore, we were surprised that blocking CaMKK starting three hours after the tLTP protocol was sufficient to prevent L-LTP expression in most experiments. Previously, we demonstrated that blocking spontaneous spiking activity with TTX from three hours to two days after tLTP induction also prevented L-LTP (*14*). Together, these results suggest that spontaneous activity in the slice culture, which is likely to be elevated specifically in FOS-expressing neurons (*46*), is essential for L-LTP expression. L-LTP requires somatic calcium elevations via L-type calcium channels (*47*).

The resulting Ca^2+^/CaM is shuttled by γCaMKII to the nucleus, where it activates CaMKK and CaMKIV (*48*). This leads to CREB activation by Ser133 phosphorylation (*49*), causing expression of FOS and other immediate early genes. In contrast, activated CaMKII in the nucleus can phosphorylate CREB at Ser142, suppressing the transcription of plasticity-related proteins (*50*). Thus, optogenetic inhibition of CaMKII may have prevented CREB phosphorylation at the inhibitory site (Fig. 8C) and enabled a robust genomic response. CaMKK activation also has local effects, it contributes to E-LTP by promoting the insertion of Ca^2+^-permeable AMPA receptors (*51*, *52*). Under conditions of CaMKII block, however, potential local effects of CaMKK were not sufficient to make L-LTP synapse-specific (Fig. 6G) and are thus not depicted in the pathway schematic (Fig. 8C).

The subsequent recruitment of PKMζ, which remains active once produced, shows that establishing L-LTP does not require an initial structural template set by CaMKII. The specificity of ZIP for PKMζ has been questioned (*53*), but we observed no effect of the scrambled control peptide, arguing against unspecific endocytosis triggered by cationic peptides at the low concentration we used (1 µM).

Contrary to models proposing that CaMKII activity is obligatory for LTP and that CaMKK functions as a downstream driver of plasticity-related protein expression (*15*), our findings demonstrate that L-LTP can be dissociated from the immediate CaMKII-dependent functional and structural changes that accompany LTP induction. However, although L-LTP was preserved in the absence of CaMKII activity, its synapse specificity was disrupted. Inhibition of CaMKII during induction resulted in a generalized potentiation of excitatory synapses onto the stimulated neurons, rather than the selective potentiation of the synapses engaged during induction. These findings from slice culture experiments raise an intriguing and testable prediction for memory function: Long-term associative memories formed under conditions of CaMKII inhibition may be comparable in strength to those formed under control conditions (*11*), yet less specific to the particular conditioned stimulus. In other words, CaMKII-dependent synaptic tagging may not be essential for establishing the final strength of an associative memory, but may instead be critical for restricting the memory trace to the synapses encoding the relevant stimulus.

Our experimental approach, optogenetic manipulation of hippocampal slice cultures, provided the stability necessary for multi-day monitoring of synaptic strength. While we do not know precisely which activity patterns activate the CaMKII-independent L-LTP pathway in the intact brain during learning, FOS expression seems to be a reliable indicator of its engagement. Furthermore, L-LTP depends on sustained neuronal activity well beyond the initial pairing protocol. This requirement for extended activity is similar to systems-level memory consolidation, during which newly acquired information is actively processed in slow-wave sleep. In vivo studies have shown that disrupting sharp-wave ripples (SWRs) and the associated replay of spike sequences impairs long-term memory formation, highlighting the functional importance of post-learning network activity (*54*). Notably, hippocampal slice cultures retain sufficient intrinsic circuitry to spontaneously generate SWRs and sequential neuronal reactivation (*55*). This preservation of endogenous network dynamics makes them a powerful and uniquely accessible model to dissect the molecular mechanisms of different phases of memory (*11*, *56*). Taken together, our findings support a model in which recurrent waves of synchronized activity, such as SWRs, engage a genomic L-LTP program through CaMKK-dependent signaling. In parallel, GluN2B-bound CaMKII acts locally at potentiated synapses to establish synaptic tags, thereby enabling the selective capture of plasticity-related proteins (*6*). In this framework, coordinated network activity and synapse-specific tagging mechanisms converge to transform transient synaptic modifications into persistent, protein synthesis–dependent forms of plasticity that support long-term memory storage.

## Acknowledgments

We thank Iris Ohmert and Jan Schröder for their excellent technical assistance, Julia Kaiser for EM data analysis. Ingke Braren from the UKE Vector Facility produced rAAVs. Xph20-eGFP (Addgene #187444) was a gift from Matthieu Sainlos, CheRiff (Addgene #51697) was a gift from Adam Cohen, ChrimsonR (Addgene #59171) was a gift from Edward Boyden, paCaMKII (Addgene #165429) was a gift from Hideji Murakoshi, LSSmOrange (Addgene #37129) was a gift from Vladislav Verkhusha, dAPEX2 (Addgene #117173) was a gift from David Ginty.

## Author contributions

Conceptualization: R.W. and T.G.O. Methodology: R.W., M.S., C.S., K.P. and T.G.O. Formal analysis: R.W., C.S. and T.G.O. Investigation: R.W. and M.S. Data curation: R.W. Writing-original draft: R.W. and T.G.O. Writing-review and editing: R.W., M.S., C.S., C.E.G., and T.G.O. Visualization: R.W. Supervision: M.S., C.E.G., and T.G.O. Project administration and funding acquisition: T.G.O. and C.E.G.

## Materials and Methods

### Animals

Sprague-Dawley rats were bred at the animal facility of the University Medical Center Hamburg-Eppendorf (UKE). All procedures were in compliance with national animal protection laws (Tierschutzgesetz der Bundesrepublik Deutschland, TierSchG). Organ harvesting was approved by the Behörde für Justiz und Verbraucherschutz (BJV) Hamburg, Lebensmittelsicherheit und Veterinärwesen (ORG1106).

#### Preparation of hippocampal slice cultures

Slice cultures were prepared from rats of either sex at postnatal day 5–7 as previously described (*57*). Briefly, hippocampi were dissected and 400 μm slices were cut from the dissected tissue with a tissue chopper. Slices were cultured on 30 mm porous membranes (Milipore PICMORG50), supplied with 1 mL of culture medium (for 500 ml: 394 ml Minimal Essential Medium (Sigma M7278), 100 ml heat inactivated donor horse serum (H1138 Sigma), 1 mM l-glutamine (Gibco 25030-024), 0.01 mg/ml insulin (Sigma I6634), 1.45 ml 5 M NaCl (S5150 Sigma), 2 mM MgSO_4_ (Fluka 63126), 1.44 mM CaCl_2_ (Fluka 21114), 0.00125% ascorbic acid (Fluka 11140), 13 mM d-glucose (Fluka 49152)) at 37°C in 5% CO_2_. The culture medium was partially (∼70%) exchanged every 3 to 5 days.

#### Transfection and viral transduction of slice cultures

Gene transfer via single-cell electroporation or viral transduction (rAAV) was performed at 10–14 days *in vitro* (DIV). To prevent non-specific activation of optogenetic tools, all procedures following transfection were conducted under dim yellow light. For sparse labeling and manipulation of CA1 pyramidal neurons, single-cell electroporation was performed as previously described (*58*). Plasmids were diluted to the desired concentration in intracellular solution containing (in mM) 135 K-gluconate, 4 MgCl_2_, 4 Na_2_-ATP, 0.4 Na-GTP, 10 Na_2_-phosphocreatine, 3 ascorbate, 10 HEPES. The following combinations were used:

- For paCaMKII two-photon imaging: pAAV-CaMP0.4-FHS-paCaMKII-WPRE3 (50 ng/μL, Addgene #165429) or paCaMKII-SD (50 ng/μL, Addgene #165430) + pCI-ZFB-Syn-Xph20-eGFP-CCR5TC (20 ng/μL, subcloned from Addgene #187444) + pCI-Syn-LSSmOrange (2 ng/μL, subcloned from Addgene #37129).
- For paCaMKII FRAP imaging: pAAV-CaMP0.4-FHS-paCaMKII-WPRE3 or paCaMKII-SD (50 ng/μL) + pCI-ZFB-Syn-Xph20-eGFP-CCR5TC (40 ng/μL) + pCI-Syn-tdimer2 (40 ng/μL).
- For paCaMKII electron tomography: pAAV-CaMP0.4-FHS-paCaMKII-WPRE3 (50 ng/μL) + pAAV-EF1α-dAPEX2 (10 ng/μL, Addgene #117173).
- For paCaMKII electrophysiology: pAAV-CaMP0.4-FHS-paCaMKII-WPRE3 or paCaMKII-SD (50 ng/μL) + pCI-syn-mKate2 (5–20 ng/μL).
- For paAIP2 electrophysiology: pAAV-hsyn-CheRiff-eGFP (0.5–2 ng/μL, Addgene #51697) + pCI-syn-mKate2 (5–20 ng/μL) + pAAV-CaMKIIP-mEGFP-P2A-paAIP2 (20 ng/μL, Addgene #91718).

For presynaptic optogenetic stimulation, a small cluster of CA3 neurons were transduced with ChrimsonR via local pressure injection of recombinant adeno-associated virus (AAV2/rh10 Syn-ChrimsonR-tdT, titer: 4.83 × 10^13^vg/mL, measured by WPRE sequence). Virus particles were diluted in HEPES buffered solution containing (in mM) 145 NaCl, 10 HEPES, 25 D-glucose, 2.5 KCl, 1 NaH_2_PO_4_, 1 MgCl_2_, 2 CaCl_2_ (pH 7.4, 314-318 mOsm/kg).

#### Chronic two-photon imaging

We equipped a two-photon microscope (Rapp OptoElectronic) with a stage incubator for multi-day recordings. The recording chamber was fabricated from titanium with a glass coverslip base and polycarbonate sidewalls. To create a semi-airtight microenvironment, the chamber top was sealed to the objective via a flexible latex membrane, and condensation was minimized using a secondary Perspex enclosure. Thermal regulation was achieved through a multi-point heating system: Two feedback-controlled heating plates positioned above and below the chamber (Okolab bold line), and two constant-current heaters attached to the water-immersion objective and oil-immersion condenser. The target temperature of 36°C was monitored by an in-bath sensor. Sterile-filtered humidified air containing 5% CO_2_ was supplied at 3 L/h (red-y compact, Vögtlin Instruments).

We scanned an area of 176 x 176 μm through a 25x/1.05 NA objective (Olympus) using ScanImage software (Vidrio) in MATLAB (R2023a). A Ti: Sapphire laser (Chameleon Ultra II, Coherent) with dispersion compensation (Spectra-Physics; Arduino Uno) was tuned to 950 nm to excite two fluorescent proteins, LSSmOrange(*20*) and GFP-labeled monobody Xph20. To correct for signal loss by scattering, laser power was exponentially adjusted with depth (Lz parameter: 350–1100). Emitted photons were captured behind objective and condenser and spectrally separated (525/50 nm and 607/70 nm filters). Z-stacks (80–150 planes, 0.75 μm spacing) were acquired (1024 × 1024 pixels) at 30 min intervals (baseline and 1.5 hours after optical stimulation) and at 60 min intervals for the rest of the recording time. Optical stimulation was delivered at 405 nm (0.1 mW/mm^2^) for 100 s (pE-4000, CoolLED).

#### Chronic two-photon image analysis

We developed an analysis pipeline (Python 3.9) consisting of denoising, rigid registration, deep-learning based segmentation and temporal tracking of detected objects (PSDs). Raw images were smoothed using a 3D median filter (r = 1 pixel) and scan artifacts (line shifts) were corrected. Time series were aligned using rigid registration based on the dendritic structure (LSSm-Orange). The PSD channel (Xph20-eGFP-CCR5TC) and the LSSM-Orange channel were used for image segmentation using star-convex object detection (StarDist) (*21*). We developed a custom napari annotation tool “SpoFi” to create ground-truth segmentation data for model training (https://github.com/drchrisch/napari-spofi). In brief, 3D images were divided into tiles (64 x 64 x 64 pixel) and 5-10 tiles were manually annotated. Model training was started from the napari plugin and predicted objects were loaded as a separate layer for visual inspection. The obtained models were retrained as necessary and then used to generate segmentation maps for the PSDs. Results were stored in a hierarchical data format (HDF5) containing raw signals, segmentation masks, and spot coordinates, paired with XML descriptors for compatibility with downstream tracking modules. Subsequently, temporal linkage of detected spines was performed using the Mastodon plugin (beta-26) in Fiji (*59*). To ensure trajectory continuity, the tracking algorithm permitted a maximum of two consecutive missing time points (gaps). Final quantitative analysis was restricted to dendritic spines that were successfully tracked across at least 80% of the recorded time frames.

For statistical analysis and plotting of data, custom data analysis scripts were written in MATLAB. To correct for photobleaching, fluorescence traces were normalized against a decay curve derived from the average intensity of 2–4 stable reference regions on the main dendritic shaft. The growth factor was defined as the relative change in fluorescence intensity over baseline. “Early Growth” refers to the mean intensity change during the 0–2 h interval post-stimulation, while “Late Growth” refers to the 2–5 h time window. For probability distribution analyses, intensity values were normalized to the specific baseline average of their parent slice to correct for inter-sample variability prior to pooling. Dendritic morphological reconstruction was performed using the SNT plugin in Fiji (*60*).

#### Measuring diffusional coupling between spine and dendrite

FRAP experiments were conducted on the same custom-built two-photon microscope as chronic imaging, using a 1070 nm fiber laser (Fidelity-2, Coherent) set to 2% power to image tdimer2. The bleach pulse was delivered to the spine head using the ‘Power Box’ function in ScanImage. 10 randomly selected spines were bleached before optical activation of paCaMKII (or paCaMKII-SD), and 10 different spines after. Exponential functions were fitted to the fluorescence recovery of every spine (Stowers plugin, Fiji). The time constants were averaged over the 10 spines and compared to the average time constant after paCaMKII activation.

#### Sample preparation for electron microscopy

Sample preparation for electron tomography followed previously established protocols (*29*), using a standard cacodylate buffer (in mM: 100 sodium cacodylate, 2 CaCl_2_, pH 7.4) for all dilutions and washing steps unless otherwise specified. About 10 minutes after paCaMKII activation (in incubator, 405nm LED, 0.1mW/mm^2^ for 100 s), slice cultures were fixed by incubation in fixation buffer (2.5% glutaraldehyde, 2% paraformaldehyde in cacodylate buffer) at 37°C for 1 min, followed by 1 h in the same fixative on ice. Samples were rinsed three times (10 min each) with ice-cold cacodylate buffer and could be stored at 4℃ for up to three weeks. To quench free aldehydes, slices were treated for 15 min with a glycine solution (in mM: 20 glycine, 100 sodium cacodylate, 2 CaCl_2_). Subsequently, endogenous peroxidase activity was blocked by incubation in freshly prepared 0.3% H_2_O_2_ (diluted from 30% stock in cacodylate buffer) for 15 min. Between steps, samples were washed 3 x in cacodylate buffer. DAB labeling was performed using a metal enhanced DAB substrate kit (Thermo Scientific, 34065); the DAB working solution was prepared immediately before use by diluting the 10x concentrate 1:9 in cacodylate buffer. Tissues were equilibrated in this solution for 1 h to ensure penetration, after which the reaction was catalyzed by adding stabilized hydrogen peroxide (1:100). Upon achieving optimal staining intensity (30-120 min), slices were washed and stabilized overnight in 3% glutaraldehyde. The following day, after micro-dissection of the CA1 region, tissues were post-fixed for 20 min on ice with 1% OsO_4_ (diluted in cacodylate buffer). Following extensive washing (1 h with exchanges every 5 min), samples were dehydrated through a graded ethanol series (30%, 50%, 70%, 80%, 90%, 100% x 2, 15 min each step) and pure propylene oxide (2 times, 10 min each). Infiltration was performed using EPON 812 resin (composition for 50 mL: 25.85 g glycidyl ether 100, 16.1 g MNA, 8.025 g DBA, 0.5 g DMP-30). The resin components were mixed at room temperature for >1 h until the color shifted from light red to golden yellow. Samples were infiltrated with resin/propylene oxide mixtures (1:1 for 90 min, 2:1 for 2 h) and then with 100% resin overnight, before curing at 58°C for 48 h.

Trimming of the block was guided by DAB-stained dendrites visualized under a Zeiss Axiophot microscope. A diamond knife was used for serial sectioning. Series of semi-thin sections (300 nm per section, 3-7 sections per grid) were collected on copper slot grids and contrasted, allowing volume reconstructions based on serial electron tomograms. The contrast staining involved 2% uranyl acetate (in 50% methanol) for 15 min, followed by lead citrate (0.04 g in 10 mL water + 5 mL 1N NaOH) for 1 min in a closed Petri dish. The lead citrate solution was centrifuged at 13000 RPM for 10 min prior to use to remove precipitates.

#### Transmission electron tomography and ultrastructural analysis

Single-axis tilt series were collected on a JEOL JEM-2100 Plus microscope operating at an acceleration voltage of 200 kV with an EMSIS CCD camera. To facilitate subsequent image alignment, 15 nm colloidal gold particles (diluted 1:20) were applied to both sides of the copper grids as fiducial markers. Regions of interest (ROIs) were selected based on positive dendritic staining patterns. For each target ROI, serial tilt series were acquired at magnifications ranging from 6000x to 10000x, covering a tilt range of approximately −60° to +60° with 1° increments, and raw data were stored in the MRC file format. Data processing involved a two-step alignment protocol: initial alignment of tilt views to generate tomograms was performed using the Etomo module of IMOD 4.11, followed by the alignment of serial tomograms for volumetric reconstruction using the TrakEM2 plugin in Fiji (*61*). Subsequent 3D reconstruction, segmentation, and visualization were carried out using Imaris 10.1.0 using the Surface_Surface_ContactArea2.py XTension (running on Python 3.7). Key structural components— including the spine head, spine neck, axon-spine interface, and postsynaptic density (PSD)—were modeled as 3D surface objects by manually tracing structure outlines. Segmentation and proofreading were performed by two experimenters both blind to the experimental conditions. Unblinding occurred only after the data analysis was complete.

#### Electrophysiology

The electrophysiology setup was based on an Olympus BX61W1 epifluorescence microscope, Axopatch 200B amplifier (Axon Instruments, Inc.), NI DAQ boards and Ephus software (in Matlab R2016b). Fiber-coupled LEDs (Mightex) and a laser combiner (Omicron LightHub) were used for multi-targeted, region-specific optical stimulation (*14*). Patch electrodes (3-4 MΩ) were filled with intracellular solution containing (in mM): 135 K-gluconate, 4 MgCl_2_, 4 Na_2_-ATP, 0.4 Na-GTP, 10 Na_2_-phosphocreatine, 3 ascorbate, and 10 HEPES (pH 7.2). The temperature was maintained at 31-33°C by heating the perfusion solution and the condenser. Recordings with series resistance (Rs) greater than 15 MΩ were excluded.

E-LTP experiments were performed at DIV 18-24 in serum-free MEM (Sigma; M7278) supplemented with (final conc. in mM) 13 D-glucose, 109 NaCl, 2 MgSO_4_, 1.44 CaCl_2_, 1 L-glutamine (Gibco 25030), 0.03 D-serine (Tocris Bioscience 0226), 0.01 mg/mL insulin (Sigma I6634), pH 7.28, 310-318 mOsm/kg. The electroporated CA1 region was centered under a 40x water immersion objective. The off-axis laser was pointed at the ChrimsonR virus-expressing CA3 region through the condenser. Regions of interest were confirmed by checking the expression of mKate2 (electroporation) or tdTomato (AAV), using an mCherry filter set to avoid unwanted activation of CheRiff, paAIP2 or paCaMKII. Recordings of the excitatory postsynaptic currents (EPSCs) were performed at −70 mV (liquid junction potential corrected) in response to light pulse stimulation of ChrimsonR-expressing CA3 neurons. In each slice culture, one CA1 cell, CheRiff-expressing or non-transfected (NT), was recorded. To obtain a baseline, ChrimsonR-expressing CA3 neurons were stimulated with 1 ms laser pulses (594 nm, 1 ms, at 0.05 Hz for 3-5 min, intensity adjusted to evoke 100-200 pA EPSCs) while EPSCs were recorded in CA1. Recordings were switched to current clamp to allow the postsynaptic neurons to spike during t-LTP induction or paCaMKII activation. The tLTP pairing protocol was one 594 nm, 2 ms light pulse through the condenser (8 mW/mm^2^) followed after 12 ms by three 405 nm, 2 ms light pulses through the objective (1.2 mW/mm^2^, 50 Hz), repeated 300 times at 5 Hz (*14*, *29*). For CaMKII inhibition experiments, paAIP2 was pre-activated for 1 s (405 nm, average power 18 μW/mm^2^) followed after 9 s by the tLTP pairing protocol. paCaMKII was activated with dim 405 nm light (100 μW/mm^2^) for 100 s. Post-stimulation EPSCs were recorded as for the baseline for 25 min. To calculate the change in input strength, the average pre-tLTP initial EPSC slope was used in equation 1 in place of the ‘average NT slope’ (Fig. S3) and the post-tLTP induction slope was used in place of the ‘TF slope’ (Fig. S3).

Intrinsic neuronal properties (Fig. S5) were characterized at DIV 18 – 27, recording in artificial cerebrospinal fluid (ACSF) containing (in mM): 119 NaCl, 11 D-glucose, 2.5 KCl, 1 NaH_2_PO_4_, 4 MgCl_2_, 4 CaCl_2_, pH 7.4, 310 mOsm/kg. Passive properties, including membrane resistance and capacitance, were derived from the response to a −5 mV test pulse. Active properties were assessed in current-clamp using a step protocol (−800 pA to +400 pA).

#### tLTP induction in the incubator

For investigating the long-term synaptic strength alteration and the expression of immediate early genes, slice cultures were optically stimulated in the incubator as previously described (*14*). Briefly, 1-2 days before stimulation, the last medium changed was performed. The slice culture insert was centered in a 35 mm culture dish and placed in a stimulation tower with two collimated high-power LEDs inside the incubator. Either tLTP induction (as described above) or paCaMKII activation (as described above) was used to induce plasticity. For immunohistochemistry, slice cultures were fixed 1 h after stimulation. For synaptic strength measurement experiments (L-LTP), slice cultures were kept in the incubator for another 1-3 days.

For pharmacological manipulation, ZIP, or scrambled ZIP (scr-ZIP), or STO-609, or TTX, was applied 3 h after incubator stimulation. ZIP and TTX were diluted to 1 µM, STO-609 (in DMSO, 5 mM stock) was diluted to 5 µM in warmed, conditioned culture medium (*14*). 50 µl of 1 µM ZIP was added directly onto the slice, followed by moving the slice to 800 µl of 1 µM ZIP. The slices were then moved back to the incubator for two days. At the end of day 2, one day before the patch-clamp readout of L-LTP, the slices were moved back to normal medium (warmed, conditioned). For quality control and data normalization, a positive (50 mM KCl) and a negative control condition (unstimulated) were included in each batch of experiments.

#### Input strength measurements

Measurements were performed between DIV 21-27 (9 to 13 days after transfection) as previously described (*14*). CheRiff-transfected CA1 neurons were patched and tested with a 1 ms blue light pulse. Neurons that did not spike in response to the light pulse were excluded from further analysis, because we could not be sure about their response to the optical tLTP protocol in the incubator. After this initial test, ChrimsonR-expressing CA3 neurons were illuminated with a 1 ms laser pulse at 594 nm, repeated 10 times at 0.05 Hz, while EPSCs were recorded in the CA1 neuron. The average of 10 sweeps was used to fit the EPSC slope. Transfected (TF) and non-transfected (NT) cells were patched in random sequence. In each slice culture, 1-5 TF cells and at least 3 NT cells were recorded and the initial slope of the EPSC (average of 6 −10 evoked responses) was determined for every cell. For each TF cell, the relative input strength was calculated as

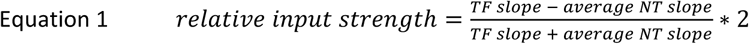

This measure of synaptic plasticity is suitable to compare variables with lognormal distributions. It is bounded between −2 and +2, with 0 = no change and 1 = three-fold increase.

To measure the input specificity of synaptic strength alterations, in a subset of experiments (Fig. 6G), an electrical stimulation electrode (filled with 500 mM NaCl) was placed close to, but not overlapping with, the ChrimsonR-expressing CA3 neurons to stimulate an independent input pathway. The stimulation current was kept constant throughout the experiment for each slice.

#### Immunohistochemistry

Slice cultures were optically stimulated to induce tLTP as described above and remained in the incubator for 1 h to allow for expression of FOS. Positive controls were stimulated 2 x for 1 minute using 50 mM KCl. Negative controls were handled as the tLTP-stimulated slices, but received no light. Drugs were diluted in conditioned culture medium to avoid the shock of fresh medium change. Slices were fixed in 4% PFA for 30 min at room temperature (RT) and then washed three times in 1x PBS for 10 min each time. To avoid unspecific primary antibody binding, slices were incubated in blocking buffer (10% (vol/vol) goat serum, 0.2% bovine serum albumin, 0.3% Triton X-100, diluted in 1x PBS) for 2 h, followed by primary antibody incubation (GFP polyclonal (Invitrogen A10262); c-Fos (Synaptic Systems 226017)) 1:1000 in carrier solution (1% (vol/vol) goat serum, 0.2% bovine serum albumin, 0.3% Triton X-100, diluted in 1x PBS) at 4℃ overnight. The next day, the slices were washed three times and incubated in secondary antibodies (Goat anti-chicken Alexa Fluor 488 (Invitrogen A11039); Goat anti-rat Alexa Fluor 647 (Invitrogen A21247); 1:1000) for another 2 h at RT. Before mounting, the slices were washed once more with either PBS or PBS containing DAPI (1:10,000). Imaging was usually performed on the same day or the first day after staining. For long-term storage, the edge of the coverslip on the mounted sections was covered with nail polish and stored in a −20 ℃ freezer. Images were acquired on a confocal microscope (Zeiss LSM900) with a 20x, 0.8NA objective. Z-stacks with 3 μm steps (1024 × 1024 pixels) were acquired for each region (around 20 to 60 μm each stack). Lasers and filters were set using the dye selection program, Zen3.5 including: DAPI, Alexa 488, Alexa 568, Alexa 647, ‘best signal’ function. All acquisitions were performed sequentially to avoid spectral cross talk. Maximum intensity projections (MIP, Fiji) were used to draw outlines of nuclei (ROIs) based on DAPI and cytoplasmic signals, blind to the FOS signal. Next, the FOS signal was quantified in each ROI (average intensity).

#### Statistical analysis

Statistical tests and sample sizes are provided in the figure legends. Results with p < 0.05 were considered statistically significant. Statistical tests were performed and data were plotted using GraphPad Prism 10 or MATLAB R2022b.

## References

1. U. Frey, R. G. Morris, Synaptic tagging and long-term potentiation. Nature 385, 533–536 (1997).

2. Y.-P. Zhang, N. Holbro, T. G. Oertner, Optical induction of plasticity at single synapses reveals input-specific accumulation of alphaCaMKII. Proc. Natl. Acad. Sci. U. S. A. 105, 12039–12044 (2008).

3. B. E. Herring, R. A. Nicoll, Long-Term Potentiation: From CaMKII to AMPA Receptor Trafficking. Annu. Rev. Physiol. 78, 351–365 (2016).

4. A. C. E. Shibata, H. H. Ueda, K. Eto, M. Onda, A. Sato, T. Ohba, J. Nabekura, H. Murakoshi, Photoactivatable CaMKII induces synaptic plasticity in single synapses. Nat. Commun. 12, 751 (2021).

5. J. Lisman, R. Yasuda, S. Raghavachari, Mechanisms of CaMKII action in long-term potentiation. Nat. Rev. Neurosci. 13, 169–182 (2012).

6. N. L. Rumian, C. M. Barker, M. E. Larsen, J. E. Tullis, R. K. Freund, A. Taslimi, S. J. Coultrap, C. L. Tucker, M. L. Dell’Acqua, K. U. Bayer, LTP expression mediated by autonomous activity of GluN2B-bound CaMKII. Cell Rep. 43, 114866 (2024).

7. J. E. Tullis, M. E. Larsen, N. L. Rumian, R. K. Freund, E. E. Boxer, C. N. Brown, S. J. Coultrap, H. Schulman, J. Aoto, M. L. Dell’Acqua, K. U. Bayer, LTP induction by structural rather than enzymatic functions of CaMKII. Nature 621, 146–153 (2023).

8. K.-U. Bayer, P. De Koninck, A. S. Leonard, J. W. Hell, H. Schulman, Interaction with the NMDA receptor locks CaMKII in an active conformation. Nature 411, 801–805 (2001).

9. E. R. Kandel, Y. Dudai, M. R. Mayford, The Molecular and Systems Biology of Memory. Cell 157, 163–186 (2014).

10. P. J. Hernandez, T. Abel, The role of protein synthesis in memory consolidation: progress amid decades of debate. Neurobiol Learn Mem 89, 293–311 (2008).

11. M. E. Shin, P. Parra-Bueno, R. Yasuda, Formation of long-term memory without short-term memory revealed by CaMKII inhibition. Nat. Neurosci. 28, 35–39 (2025).

12. A. Goto, A. Bota, K. Miya, J. Wang, S. Tsukamoto, X. Jiang, D. Hirai, M. Murayama, T. Matsuda, T. J. McHugh, T. Nagai, Y. Hayashi, Stepwise synaptic plasticity events drive the early phase of memory consolidation. Science 374, 857–863 (2021).

13. H. Murakoshi, M. E. Shin, P. Parra-Bueno, E. M. Szatmari, A. C. E. Shibata, R. Yasuda, Kinetics of Endogenous CaMKII Required for Synaptic Plasticity Revealed by Optogenetic Kinase Inhibitor. Neuron 94, 37–47.e5 (2017).

14. M. Anisimova, B. van Bommel, R. Wang, M. Mikhaylova, J. S. Wiegert, T. G. Oertner, C. E. Gee, Spike-timing-dependent plasticity rewards synchrony rather than causality. Cereb. Cortex 33, 23–34 (2022).

15. R. L. Redondo, H. Okuno, P. A. Spooner, B. G. Frenguelli, H. Bito, R. G. M. Morris, Synaptic Tagging and Capture: Differential Role of Distinct Calcium/Calmodulin Kinases in Protein Synthesis-Dependent Long-Term Potentiation. J. Neurosci. 30, 4981–4989 (2010).

16. H. Patel, R. Zamani, The role of PKMζ in the maintenance of long-term memory: a review. Reviews in the Neurosciences 32, 481–494 (2021).

17. T. C. Sacktor, A. A. Fenton, What does LTP tell us about the roles of CaMKII and PKMζ in memory? Mol Brain 11, 77 (2018).

18. G. Girardeau, K. Benchenane, S. I. Wiener, G. Buzsáki, M. B. Zugaro, Selective suppression of hippocampal ripples impairs spatial memory. Nat. Neurosci. 12, 1222–1223 (2009).

19. C. Rimbault, C. Breillat, B. Compans, E. Toulmé, F. N. Vicente, M. Fernandez-Monreal, P. Mascalchi, C. Genuer, V. Puente-Muñoz, I. Gauthereau, E. Hosy, S. Claverol, G. Giannone, I. Chamma, C. D. Mackereth, C. Poujol, D. Choquet, M. Sainlos, Engineering paralog-specific PSD-95 recombinant binders as minimally interfering multimodal probes for advanced imaging techniques. Elife 13 (2024).

20. D. M. Shcherbakova, M. A. Hink, L. Joosen, T. W. J. Gadella, V. V. Verkhusha, An Orange Fluorescent Protein with a Large Stokes Shift for Single-Excitation Multicolor FCCS and FRET Imaging. J. Am. Chem. Soc. 134, 7913–7923 (2012).

21. M. Weigert, U. Schmidt, R. Haase, K. Sugawara, G. Myers, “Star-convex Polyhedra for 3D Object Detection and Segmentation in Microscopy” in 2020 IEEE Winter Conference on Applications of Computer Vision (WACV) (2020), pp. 3655–3662.

22. Y. Loewenstein, A. Kuras, S. Rumpel, Multiplicative Dynamics Underlie the Emergence of the Log-Normal Distribution of Spine Sizes in the Neocortex In Vivo. Journal of Neuroscience 31, 9481–9488 (2011).

23. J. Karbowski, P. Urban, Information encoded in volumes and areas of dendritic spines is nearly maximal across mammalian brains. Sci Rep 13, 22207 (2023).

24. L. Hazan, N. E. Ziv, Activity Dependent and Independent Determinants of Synaptic Size Diversity. J Neurosci 40, 2828–2848 (2020).

25. M. F. Eggl, T. E. Chater, J. Petkovic, Y. Goda, T. Tchumatchenko, Linking spontaneous and stimulated spine dynamics. Commun Biol 6, 1–13 (2023).

26. J. I. Arellano, R. Benavides-Piccione, J. Defelipe, R. Yuste, Ultrastructure of dendritic spines: correlation between synaptic and spine morphologies. Front Neurosci 1, 131–143 (2007).

27. J. Tønnesen, G. Katona, B. Rózsa, U. V. Nägerl, Spine neck plasticity regulates compartmentalization of synapses. Nat Neurosci 17, 678–685 (2014).

28. Q. Zhang, W.-C. A. Lee, D. L. Paul, D. D. Ginty, Multiplexed peroxidase-based electron microscopy labeling enables simultaneous visualization of multiple cell types. Nat. Neurosci. 22, 828–839 (2019).

29. R. Wang, M. Schweizer, M. Anisimova, C. E. Gee, T. G. Oertner, Ultrastructural analysis of synapses after induction of spike-timing-dependent plasticity. *Cell Rep*. Methods 5, 101142 (2025).

30. R. Araya, T. P. Vogels, R. Yuste, Activity-dependent dendritic spine neck changes are correlated with synaptic strength. Proceedings of the National Academy of Sciences 111, E2895–E2904 (2014).

31. M. Bosch, J. Castro, T. Saneyoshi, H. Matsuno, M. Sur, Y. Hayashi, Structural and molecular remodeling of dendritic spine substructures during long-term potentiation. Neuron 82, 444–459 (2014).

32. A. Grunditz, N. Holbro, L. Tian, Y. Zuo, T. G. Oertner, Spine neck plasticity controls postsynaptic calcium signals through electrical compartmentalization. J. Neurosci. 28, 13457–13466 (2008).

33. B. L. Bloodgood, B. L. Sabatini, Neuronal Activity Regulates Diffusion Across the Neck of Dendritic Spines. Science 310, 866–869 (2005).

34. A. M. Shirke, R. Malinow, Mechanisms of potentiation by calcium-calmodulin kinase II of postsynaptic sensitivity in rat hippocampal CA1 neurons. J Neurophysiol 78, 2682–2692 (1997).

35. J. Jaworski, K. Kalita, E. Knapska, c-Fos and neuronal plasticity: the aftermath of Kaczmarek’s theory. Acta Neurobiol Exp (Wars) 78, 287–296 (2018).

36. C. M. Alberini, Transcription factors in long-term memory and synaptic plasticity. Physiol. Rev. 89, 121–145 (2009).

37. N. Caporale, Y. Dan, Spike Timing–Dependent Plasticity: A Hebbian Learning Rule. Annual Review of Neuroscience 31, 25–46 (2008).

38. K. Barcomb, J. W. Hell, T. A. Benke, K. U. Bayer, The CaMKII/GluN2B Protein Interaction Maintains Synaptic Strength*. Journal of Biological Chemistry 291, 16082–16089 (2016).

39. W. Tao, J. Lee, X. Chen, J. Díaz-Alonso, J. Zhou, S. Pleasure, R. A. Nicoll, Synaptic memory requires CaMKII. Elife 10, e60360 (2021).

40. M. Sanhueza, G. Fernandez-Villalobos, I. S. Stein, G. Kasumova, P. Zhang, K. U. Bayer, N. Otmakhov, J. W. Hell, J. Lisman, Role of the CaMKII/NMDA receptor complex in the maintenance of synaptic strength. J. Neurosci. 31, 9170–9178 (2011).

41. M. J. LeBlancq, T. L. McKinney, C. T. Dickson, ZIP It: Neural Silencing Is an Additional Effect of the PKM-Zeta Inhibitor Zeta-Inhibitory Peptide. J. Neurosci. 36, 6193–6198 (2016).

42. N. Sadeh, S. Verbitsky, Y. Dudai, M. Segal, Zeta Inhibitory Peptide, a Candidate Inhibitor of Protein Kinase M, Is Excitotoxic to Cultured Hippocampal Neurons. Journal of Neuroscience 35, 12404–12411 (2015).

43. R. L. Redondo, R. G. M. Morris, Making memories last: the synaptic tagging and capture hypothesis. Nat Rev Neurosci 12, 17–30 (2011).

44. C. P. Goold, R. A. Nicoll, Single-cell optogenetic excitation drives homeostatic synaptic depression. Neuron 68, 512–528 (2010).

45. C. N. Brown, K. U. Bayer, Studying CaMKII: Tools and standards. Cell Reports 43 (2024).

46. M. Pignatelli, T. J. Ryan, D. S. Roy, C. Lovett, L. M. Smith, S. Muralidhar, S. Tonegawa, Engram Cell Excitability State Determines the Efficacy of Memory Retrieval. Neuron 101, 274–284.e5 (2019).

47. C. R. Raymond, S. J. Redman, Spatial segregation of neuronal calcium signals encodes different forms of LTP in rat hippocampus. J. Physiol. 570, 97–111 (2006).

48. H. Ma, R. D. Groth, S. M. Cohen, J. F. Emery, B. Li, E. Hoedt, G. Zhang, T. A. Neubert, R. W. Tsien, γCaMKII shuttles Ca^2+^/CaM to the nucleus to trigger CREB phosphorylation and gene expression. Cell 159, 281–294 (2014).

49. G. Y. Wu, K. Deisseroth, R. W. Tsien, Activity-dependent CREB phosphorylation: convergence of a fast, sensitive calmodulin kinase pathway and a slow, less sensitive mitogen-activated protein kinase pathway. Proc Natl Acad Sci U S A 98, 2808–2813 (2001).

50. P. Sun, H. Enslen, P. S. Myung, R. A. Maurer, Differential activation of CREB by Ca2+/calmodulin-dependent protein kinases type II and type IV involves phosphorylation of a site that negatively regulates activity. Genes Dev. 8, 2527–2539 (1994).

51. E. S. Guire, M. C. Oh, T. R. Soderling, V. A. Derkach, Recruitment of calcium-permeable AMPA receptors during synaptic potentiation is regulated by CaM-kinase I. J. Neurosci. 28, 6000–6009 (2008).

52. D. A. Fortin, M. A. Davare, T. Srivastava, J. D. Brady, S. Nygaard, V. A. Derkach, T. R. Soderling, Long-term potentiation-dependent spine enlargement requires synaptic Ca2+-permeable AMPA receptors recruited by CaM-kinase I. J. Neurosci. 30, 11565–11575 (2010).

53. E. G. Stokes, J. J. Vasquez, G. Azouz, M. Nguyen, A. Tierno, Y. Zhuang, V. M. Galinato, M. Hui, M. Toledano, I. Tyler, X. Shi, R. F. Hunt, J. Aoto, K. T. Beier, Cationic peptides cause memory loss through endophilin-mediated endocytosis. Nature 638, 479–489 (2025).

54. G. M. van de Ven, S. Trouche, C. G. McNamara, K. Allen, D. Dupret, Hippocampal offline reactivation consolidates recently formed cell assembly patterns during sharp wave-ripples. Neuron 92, 968–974 (2016).

55. T. Ishikawa, Y. Ikegaya, Locally sequential synaptic reactivation during hippocampal ripples. Sci. Adv. 6, eaay1492 (2020).

56. S. Zhai, E. D. Ark, P. Parra-Bueno, R. Yasuda, Long-distance integration of nuclear ERK signaling triggered by activation of a few dendritic spines. Science 342, 1107–1111 (2013).

57. C. E. Gee, I. Ohmert, J. S. Wiegert, T. G. Oertner, Preparation of Slice Cultures from Rodent Hippocampus. Cold Spring Harb. Protoc. 2017 (2017).

58. J. S. Wiegert, C. E. Gee, T. G. Oertner, Single-cell electroporation of neurons. Cold Spring Harb. Protoc. 2017, db.prot094904 (2017).

59. C. Wolff, J.-Y. Tinevez, T. Pietzsch, E. Stamataki, B. Harich, L. Guignard, S. Preibisch, S. Shorte, P. J. Keller, P. Tomancak, A. Pavlopoulos, Multi-view light-sheet imaging and tracking with the MaMuT software reveals the cell lineage of a direct developing arthropod limb. Elife 7, e34410 (2018).

60. C. Arshadi, U. Günther, M. Eddison, K. I. S. Harrington, T. A. Ferreira, SNT: a unifying toolbox for quantification of neuronal anatomy. Nat Methods 18, 374–377 (2021).

61. A. Cardona, S. Saalfeld, J. Schindelin, I. Arganda-Carreras, S. Preibisch, M. Longair, P. Tomancak, V. Hartenstein, R. J. Douglas, TrakEM2 Software for Neural Circuit Reconstruction. PLOS ONE 7, e38011 (2012).

