## Supplemental Figures 1-9 for "Dissecting the molecular triggers of early and late long-term potentiation"

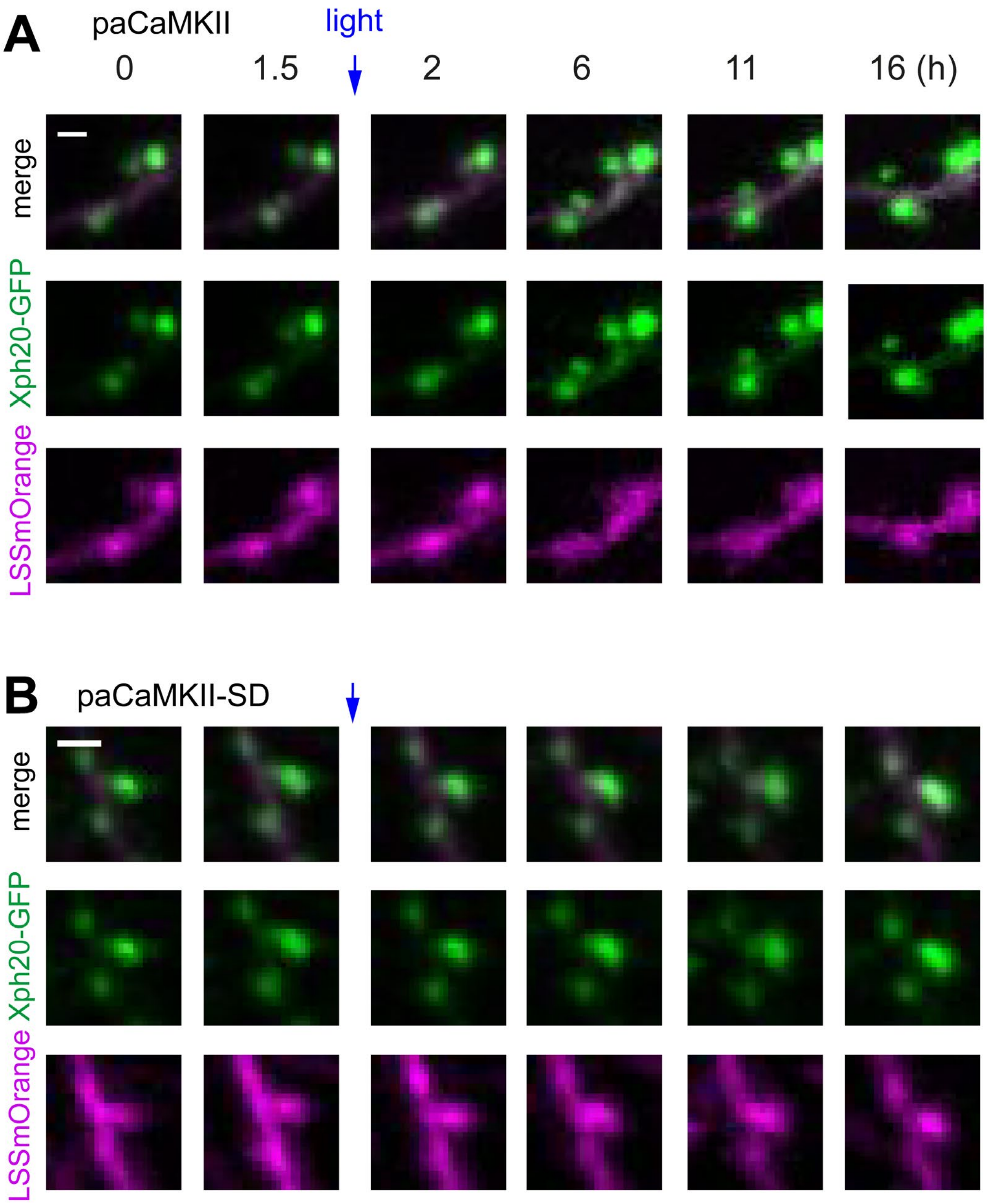


**Fig. S1. Tracking PSD and spine volume over time.**

**(A)** Enlarged detail of an imaging experiment with paCaMKII activation (blue arrow). Green: PSD monobody Xph20-GFP. Magenta: volume-filling LSSmOrange. **(B)** Example images from a control experiment (paCaMKII-SD). Scale bars: 1.5 µm.

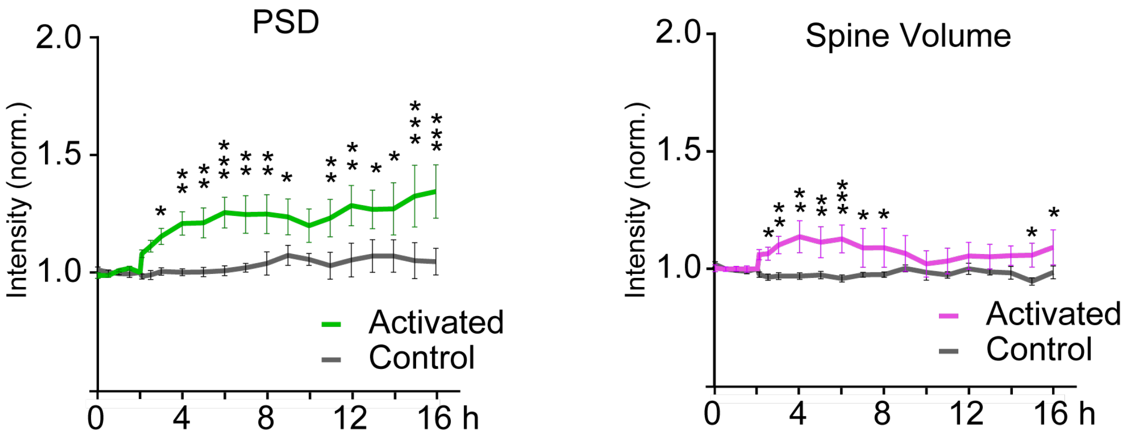


| | |
|---|---|
| Mixed-effects model (REML). Time, F (20, 219) = 6.484, P < 0.0001; optical stimulation, F (1, 14) = 12.09, P = 0.0037; Interaction, F (20, 219) = 3.281, P < 0.0001. | Šídák's, multiple comparisons P values from baseline (x4) to post stimulation effects (x16): 0.701, 0.8283, 0.829, 0.7003, 0.1278, 0.0551, 0.0195, 0.0018, 0.0016, 0.0002, 0.0011, 0.0031, 0.0198, 0.0704, 0.0098, 0.003, 0.0115, 0.0104, 0.0004, 0.0001 |

| | |
|---|---|
| Mixed-effects model (REML). Time, F (20, 225) = 5.428, P < 0.0001; optical stimulation, F (1, 14) = 3.063, P = 0.1020; Interaction, F (20, 225) = 1.490, P = 0.0861. | Šídák's, multiple comparisons P values from baseline (x4) to post stimulation effects (x16): 0.8389, 0.9229, 0.8848, 0.8767, 0.8377, 0.7565, 0.5832, 0.3419, 0.3094, 0.1662, 0.0655, 0.0617, 0.0586, 0.0823, 0.0454, 0.0156, 0.0544, 0.1106, 0.037, 0.0162. |

**Fig. S2. Statistical test of individual time points after activation of paCaMKII**

Same data as Fig. 1E. Data plotted as mean ± SEM. Statistical test methods and results are listed under the plot.

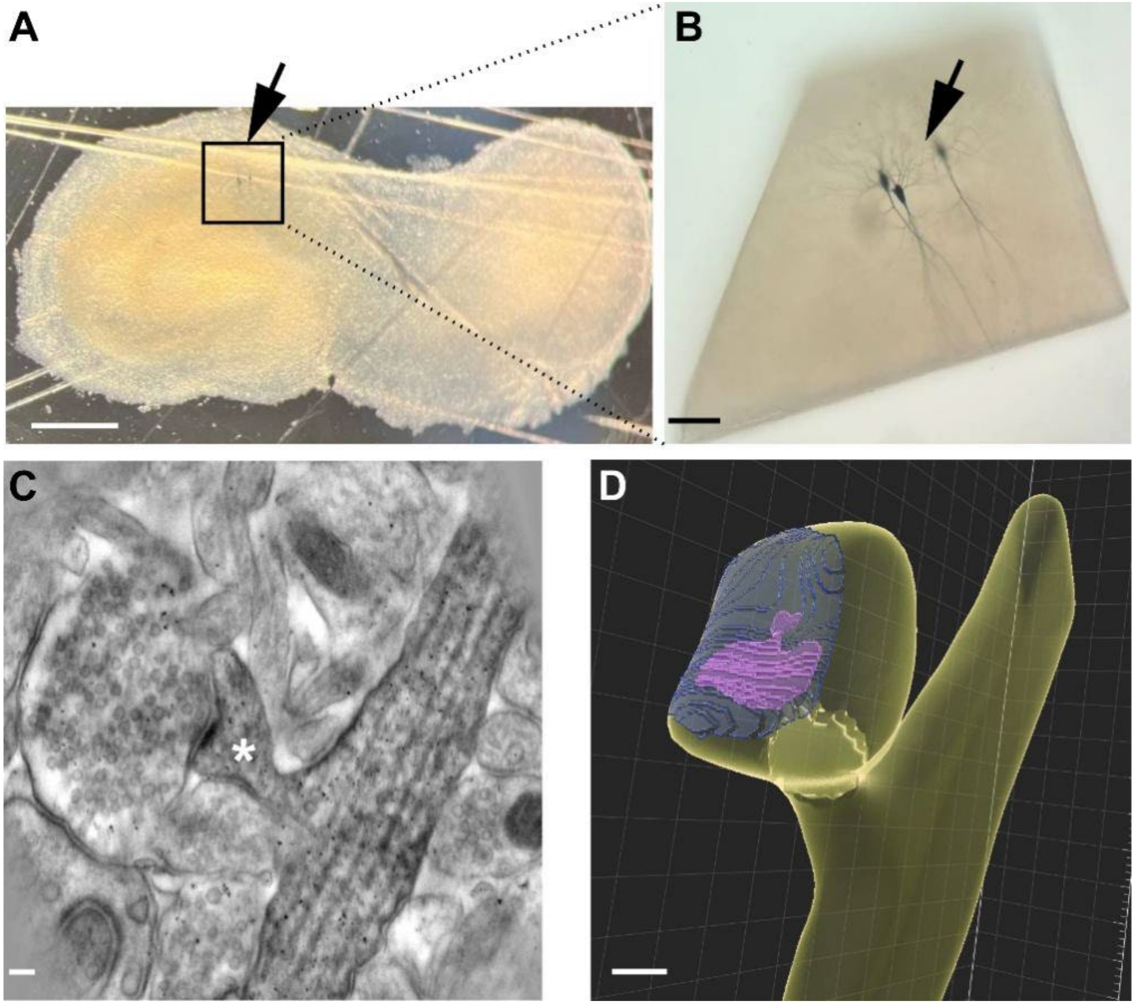


**Fig. S3. dAPEX2 labeling of spines after paCaMKII activation.**

**(A)** After DAB staining, transfected PNs appeared dark-brown (arrow). Scale bar: 700 µm. **(B)** Stained cells were dissected for sectioning. Scale bar: 60 µm. **(C)** Reconstruction of labeled dendrite and spine (white asterisk) from tilt series. Scale bar: 100 nm. **(D)** 3D surface view (Imaris) of the labeled spine in the EM image. Yellow: spine head and parent dendrite; magenta: PSD; transparent blue: axon-spine interface. Scale bar: 100 nm.

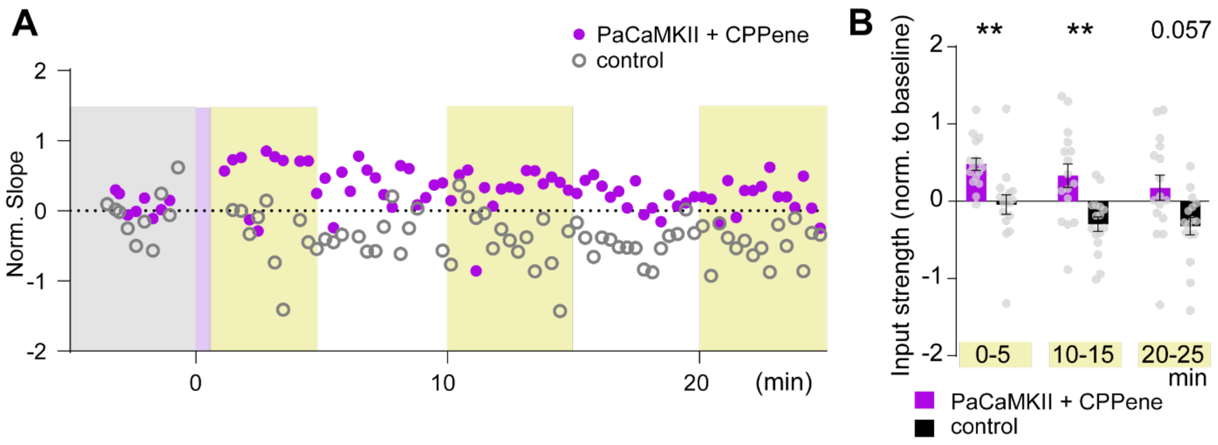


**Fig. S4. CaMKII activation-induced input strengthening does not require NMDAR activity.** **(A)** Whole-cell patch clamp recordings of synaptic strength upon paCaMKII activation at t = 0 (violet bar) under conditions of NMDA receptor block (CPPene). Control neuron expresses only the fluorescent protein mKate2 (gray rings). **(B)** Summary of input strength changes after light stimulation in three time windows (yellow shading). CPPene-paCaMKII: n = 16 slice cultures. Control: n = 16 slice cultures. Two-way ANOVA followed by Dunnett's multiple comparisons. Data plotted as mean ± SEM. **$p < 0.01$.

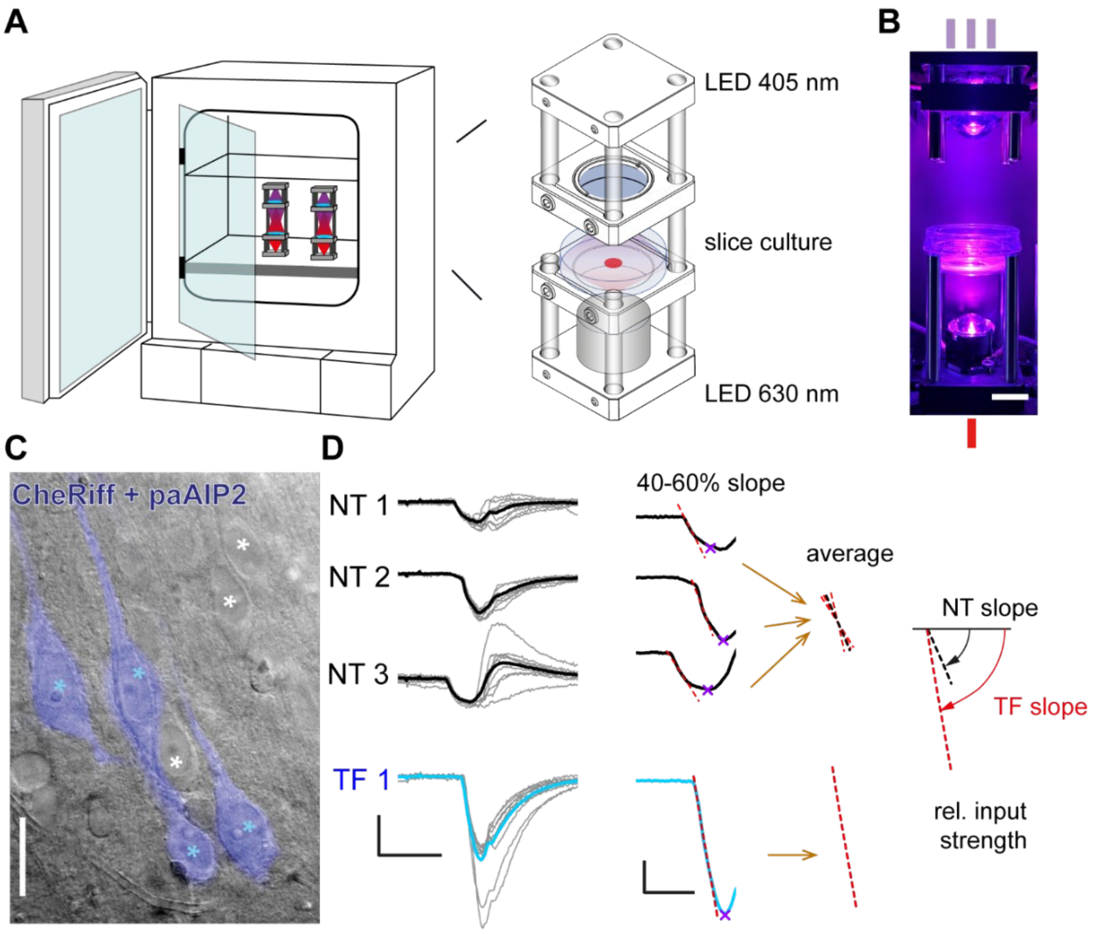


**Fig. S5. Relative input strength calculation 1 to 3 days after incubator optical stimulation.** **(A)** The setup of the incubator stimulation. Left: Stimulation towers placed inside the culture incubator. Right: Optical stimulation was delivered by collimated LED light sources, controlled from outside the incubator. Figures adapted from Anisimova et al., 2022. Slices were kept in the incubator for 1 to 3 days after optical stimulation, before the input strength was determined in patch-camp experiments. **(B)** The LED stimulation tower with a 35 mm Petri dish containing the cell culture insert. Ticks symbolize the three 405 nm light pulses from the top LED and the single 635 nm pulse from the bottom LED. **(C)** Dodt contrast image of the CA1 region with overlaid epifluorescence image of transfected (TF, blue asterisk) pyramidal neurons (in this example expressing CheRiff and mKate2). Neighboring unlabeled cells are non-transfected neurons (NT, white asterisks). Scale bar: 36 µm. **(D)** Read-out of input strength. EPSCs were recorded from NT and TF CA1 pyramidal neurons (thin grey lines) and averaged (thick black/blue lines, average of 8 - 10 sweeps). The first peak of the average EPSC was detected (magenta cross) and the initial slope measured (red dashed lines). The slope values of NT cells were averaged and the relative input strength was calculated: rel. input strength = 2*(TF slope – average NT slope)/ (TF slope + average NT slope). The intensity of the laser was kept constant for all recordings from the same slice. PNs were sequentially recorded from a single field of view per slice without stage movements to ensure consistent laser stimulation of inputs for all recorded neurons. Scale bars: 100 ms, 50 pA (left), 50 ms, 25 pA (right).

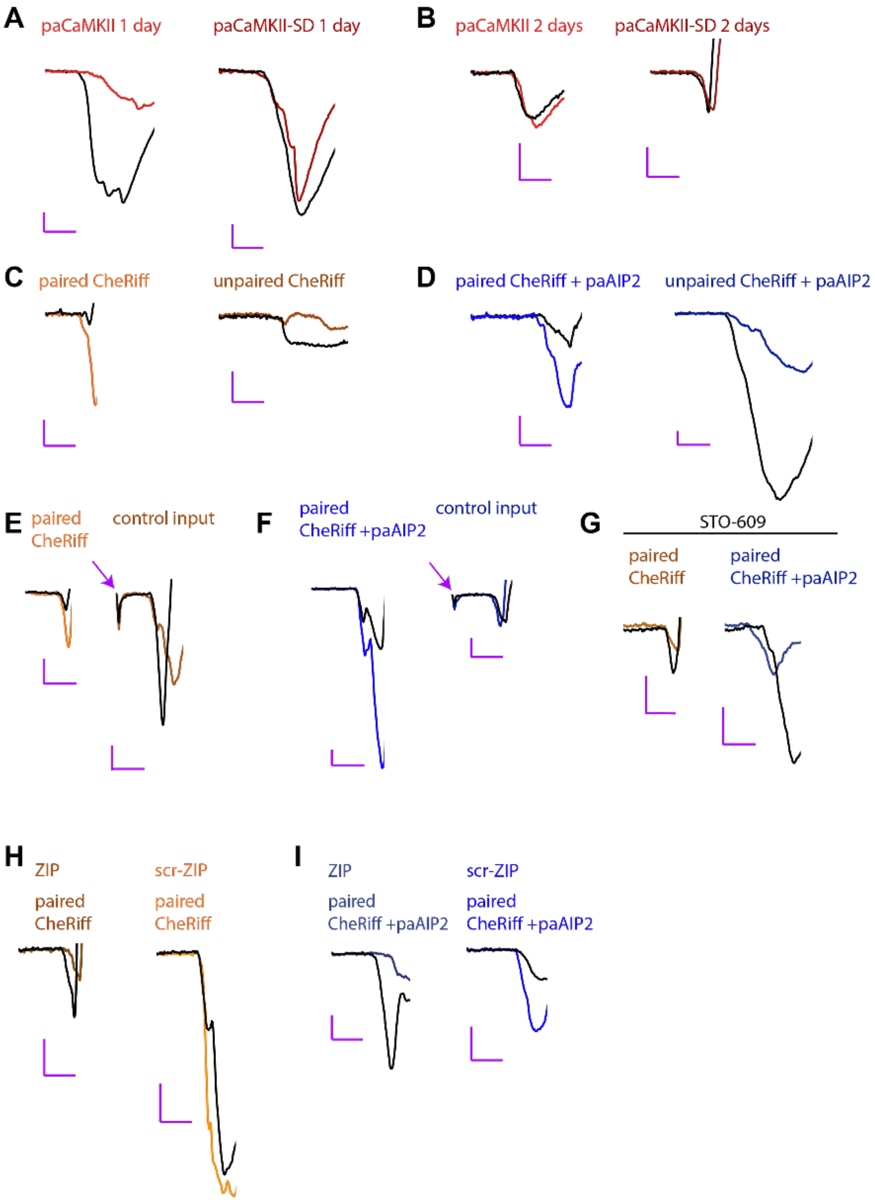


**Fig. S6. Example traces recorded in CA1 neurons in response to optogenetic activation of CA3 neurons.**

Colored lines are EPSCs from transfected CA1 pyramidal cells, black lines are EPSCs in neighboring (non-transfected) pyramidal cells, evoked by the same optogenetic stimulation of CA3. Note that slope measurements and input strength calculation were performed on averages (see Fig. S5), not on single trial responses as shown here. (**A-B**) examples for Fig. 4D, paCaMKII and paCaMKII-SD input strength 1 or 2 days after incubator stimulation. (**C-D**) examples for Fig. 5F, optically paired and unpaired CheRiff cells or paAIP2 + CheRiff cells input strength 3 days after incubator pairing. (**E-F**) examples for Fig. 5G, optically evoked and electrically evoked CheRiff or CheRiff + paAIP2 cells input strength 3 days after incubator optical pairing. Arrows: stimulation artifact. (**G-I**) examples for Fig. 8B, paired CheRiff and CheRiff + paAIP2 cells, with STO-609, ZIP or scr-ZIP incubation, input strength 3 days after in-incubator optical pairing.

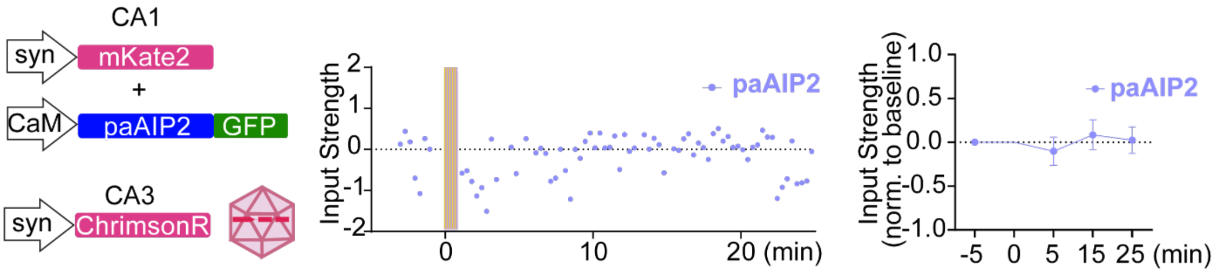


**Fig. S7. Expression and activation of paAIP2 does not affect basal transmission.**

CA3 cells were virally transduced with ChrimsonR and CA1 cells electroporated with paAIP2 and mKate2. Example recording shows that CaMKII inhibition by optical activation of paAIP2 does not affect synaptic strength within 25 min. Right: Average of n = 10 experiments, normalized to baseline (mean ± SEM).

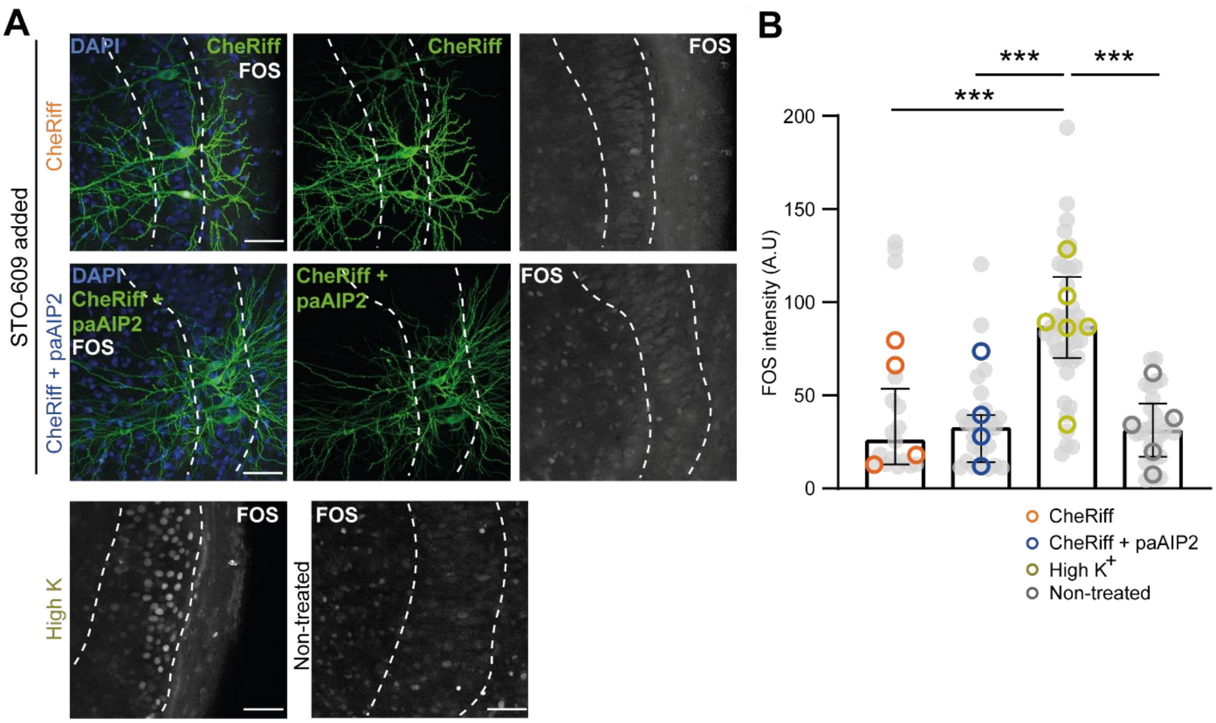


**Fig. S8. CaMKK is critical for FOS expression.**

**(A)** Optical tLTP protocol applied under conditions of CaMKK block (STO-609). Slice cultures were incubated in STO-609 for 12-24 h before tLTP induction. Cell body layer is indicated by dashed lines. Slice cultures were fixed 1 h after tLTP induction. Anti-FOS immunoreactivity is shown in white. Scale bar: 50 µm. **(B)** Evaluation of FOS immunoreactivity. With ('CheRiff') or without ('CheRiff + paAIP2') CaMKII activity, blocking CaMKK ('STO-609 added') prevents FOS expression after optical pairing, with FOS expression levels significantly different from positive (high $K^+$) controls. Kruskal-Wallis test followed by Dunn's multiple comparisons test, ***$p < 0.001$, n = 17, 26,45, 29 cells (grey dots); bars show median ± quartiles. Open circles: Repeats (N = 4, 4, 6, 5 slice cultures). A.U., arbitrary units.

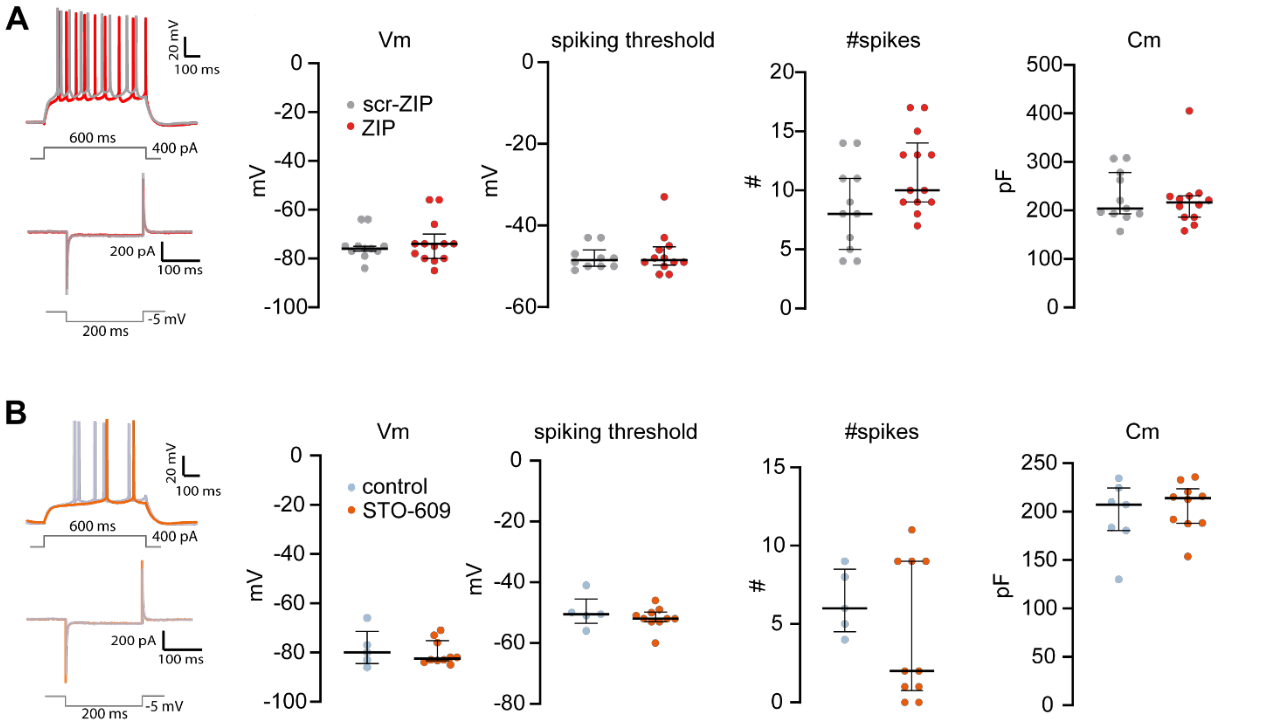


**Fig. S9. Neither ZIP nor STO-609 affected cell parameters.**

**(A)** Example traces of spike trains evoked by 400 pA current injections (red: ZIP; gray: scrZIP) and -5 mV test pulse response (voltage clamp). Bath application of ZIP did not have significant effects on resting membrane voltage (Vm), spiking threshold, number of spikes evoked by 400 pA current injections, or membrane capacitance (Cm). Control neurons expressed scrambled ZIP (scr-ZIP, light gray). Data plotted as median ± IQR. From left to right, n = 11, 13; 10, 12; 11, 13; 11, 12 neurons. Mann-Whitney test. **(B)** Bath application of STO-609 did not have significant effects on resting membrane voltage (Vm), spiking threshold, number of spikes evoked by 400 pA current injections, or membrane capacitance (Cm). Data plotted as median ± IQR. From left to right, n = 5, 10; 5, 10; 5, 10; 7, 10 slice cultures. Mann-Whitney tests.
